# Effect of SARS-CoV-2 spike mutations on animal ACE2 usage and in vitro neutralization sensitivity

**DOI:** 10.1101/2021.01.27.428353

**Authors:** Weitong Yao, Danting Ma, Haimin Wang, Xiaojuan Tang, Chengzhi Du, Hong Pan, Chao Li, Hua Lin, Michael Farzan, Jincun Zhao, Yujun Li, Guocai Zhong

**Affiliations:** School of Chemical Biology and Biotechnology, Peking University Shenzhen Graduate School, Shenzhen 518055, China; Shenzhen Bay Laboratory, Shenzhen 518132, China; Biomedical Research Center of South China, Fujian Normal University, Fuzhou 350117, China; Department of Immunology and Microbiology, The Scripps Research Institute, Jupiter, Florida 33458, United States; State Key Laboratory of Respiratory Disease, National Clinical Research Center for Respiratory Disease, Guangzhou Institute of Respiratory Health, the First Affiliated Hospital of Guangzhou Medical University, Guangzhou, Guangdong, China 510120

**Keywords:** SARS-CoV-2, spike mutation, variant of concern, receptor, ACE2, host range, neutralizing antibody, ACE2-Ig

## Abstract

The emergence of SARS-CoV-2 variants poses greater challenges to the control of COVID-19 pandemic. Here, we parallelly investigated three important characteristics of seven SARS-CoV-2 variants, including two mink-associated variants, the B.1.617.1 variant, and the four WHO-designated variants of concerns (B.1.1.7, B.1.351, P.1, and B.1.617.2). We first investigated the ability of these variants to bind and use animal ACE2 orthologs as entry receptor. We found that, in contrast to a prototype variant, the B.1.1.7, B.1.351, and P.1 variants had significantly enhanced affinities to cattle, pig, and mouse ACE2 proteins, suggesting increased susceptibility of these species to these SARS-CoV-2 variants. We then evaluated in vitro neutralization sensitivity of these variants to four monoclonal antibodies in clinical use. We observed that all the variants were partially or completely resistant against at least one of the four tested antibodies, with B.1.351 and P.1 showing significant resistance to three of them. As ACE2-Ig is a broad-spectrum anti-SARS-CoV-2 drug candidate, we then evaluated in vitro neutralization sensitivity of these variants to eight ACE2-Ig constructs previously described in three different studies. All the SARS-CoV-2 variants were efficiently neutralized by these ACE2-Ig constructs. Interestingly, compared to the prototype variant, most tested variants including the variants of concern B.1.1.7, B.1.351, P.1, and B.1.617.2 showed significantly increased (up to ∼15-fold) neutralization sensitivity to ACE2-Ig constructs that are not heavily mutated in the spike-binding interface of the soluble ACE2 domain, suggesting that SARS-CoV-2 evolves toward better utilizing ACE2, and that ACE2-Ig is an attractive drug candidate for coping with SARS-CoV-2 mutations.

## Introduction

As of June 23^rd^ 2021, the severe acute respiratory syndrome coronavirus 2 (SARS-CoV-2), the etiological agent of the ongoing coronavirus disease 2019 (COVID-19) pandemic, has already caused about 178 million confirmed infections and over 3.8 million documented deaths over the world, according to World Health Organization (WHO) online updates. The pandemic has triggered unprecedentedly extensive worldwide efforts to develop countermeasures against COVID-19, and a number of encouraging progresses have been achieved in developing prophylactic vaccines and antibody therapeutics^1-12^. So far, there are more than six prophylactic COVID-19 vaccines that have been authorized by different countries for emergency use. These include two mRNA vaccines (Pfizer-BioNTech, US; Moderna, US)^1,2^, two inactivated vaccines (Sinopharm, China; Sinovac, China)^3-5^, and two adenoviral vectored vaccines (Sputnik V, Russia; AstraZeneca-Oxford, UK)^6,7^. Over 2 billion doses of these vaccines have been administered worldwide. There are also some convalescent patient-derived antibodies that have been authorized for emergency use by the US FDA, such as Regeneron’s antibody cocktail consisting of casirivimab (REGN10933) and imdevimab (REGN10987)^8,9^, and Eli Lilly’s antibody cocktail consisting of etesevimab (LY-CoV016) and bamlanivimab (LY-CoV555)^10-12^. So far, all the SARS-CoV-2 vaccines and antibody therapeutics in clinical use were developed on the basis of the prototype SARS-CoV-2 strain.

SARS-CoV-2 is a betacoronavirus that has broad host ranges^13-16^. Receptor usage is a critical determinant for the host range, as well as an effective neutralization target, of coronaviruses ^16,17^. SARS-CoV-2 utilizes ACE2 as a key cellular receptor to infect cells^13,18,19^. Antibodies targeting the interactions between ACE2 and SARS-CoV-2 spike receptor-binding domain (RBD) efficiently neutralize SARS-CoV-2 infection and reduced viral load in animal models and COVID-19 patients^8-12^. As human ACE2 residues on the spike RBD-binding interface are highly conserved across a number of mammalian-ACE2 orthologs, mutations within the spike RBD region might easily alter cross-species receptor usage by SARS-CoV-2, as well as sensitivity to SARS-CoV-2 neutralization agents. Indeed, we recently found that SARS-CoV-2 can use human ACE2 and a wide range of animal-ACE2 orthologs, but not mouse ACE2, for cell entry ^14^. But a single amino-acid change within the spike receptor-binding domain (RBD; Q498H, Q498Y, or N501Y) could be sufficient to confer SARS-CoV-2 the ability to utilize mouse ACE2^ref.20-23^.

SARS-CoV-2 is a single-stranded RNA virus with moderate mutation and recombination frequencies^24,25^. A number of spontaneous and selection-pressure-driven mutations of SARS-CoV-2 genome have been identified in viral variants that emerged during the course of the pandemic^13,26-35^. With more and more SARS-CoV-2 variants being identified to carry diverse spike mutations within the RBD region, it’s possible that some of the spike mutations might alter the host range of the virus, or compromise the efficacy of vaccines or neutralizing antibodies developed on the basis of the prototype SARS-CoV-2 strain. In this study, we investigated cross-species receptor usage of multiple circulating SARS-CoV-2 variants that emerged during the pandemic, as well as sensitivity of these variants to four neutralizing antibodies in clinical use and a broad-spectrum anti-SARS-CoV-2 drug candidate.

## Results

### Spike variants and related mutations investigated in this study

We first cloned twelve spike genes of nine SARS-CoV-2 natural variants. They include a spike gene of an early isolate WHU01^ref.36^, representing the prototype SARS-CoV-2 variant, a spike gene of the well-studied D614G variant^35^, two spike genes of a mink-associated variant that carries a Y453F signature mutation in the RBD region^31,32,38^, a spike gene of the B.1.617.1 variant, and seven spike genes of the four WHO-designated variants of concern (VOCs), B.1.1.7, B.1.351, P.1, and B.1.617.2^ref.27-30^ (**Figure 1A**). The VOC B.1.1.7, also called 501Y.V1, was first identified in the UK in late summer to early autumn 2020, then spread rapidly across the UK, and now has been identified in at least 114 countries^39-41^. B.1.1.7 has an N501Y signature mutation in the RBD region, and carries seven additional mutations in the remaining region of the spike protein^27,28^. Two spike sequences, one that only carries the N501Y mutation and the other that carries all the spike mutations, were used in the following studies (**Figure 1A**). The VOC B.1.351, also called 501Y.V2, was first detected in South Africa from samples collected in early August, then rapidly became dominant locally, and now has been identified in at least 68 countries. B.1.351 variant has three signature amino-acid substitutions (K417N-E484K-N501Y) in the RBD region and a number of additional mutations in the remaining regions of the spike protein^29^. Similarly, two spike sequences, one that only carries the K417N-E484K-N501Y mutations and the other that carries all the spike mutations, were used in the following studies (**Figure 1A**). The VOC P.1, also called 501Y.V3, emerged in Brazil around mid-November 2020, and now has rapidly spread to at least 37 countries. P.1 variant has three signature substitutions (K417T-E484K-N501Y) in the RBD region, similar to the RBD mutations of the B.1.351 variant. It also carries seven additional lineage-defining amino-acid substitutions in the remaining regions of the spike protein^30^. Again, two spike sequences, one that only carries the K417T-E484K-N501Y mutations and the other that carries all the spike mutations, were used in the following studies (**Figure 1A**). The VOC B.1.617.2, emerged in India around September 2020, and now has rapidly spread to over 75 countries. B.1.617.2 variant has two signature substitutions (L452R-T478K) in the RBD region. It also carries six additional lineage-defining amino-acid substitutions in the remaining regions of the spike protein. A spike sequence that carries all the B.1.617.2-signature mutations was used in the following studies (**Figure 1A**). All the residues associated with the above mentioned RBD mutations are indicated in the structure of human ACE2 in complex with the RBD of a prototype SARS-CoV-2 variant (**Figure 1B**).

**Figure 1.**
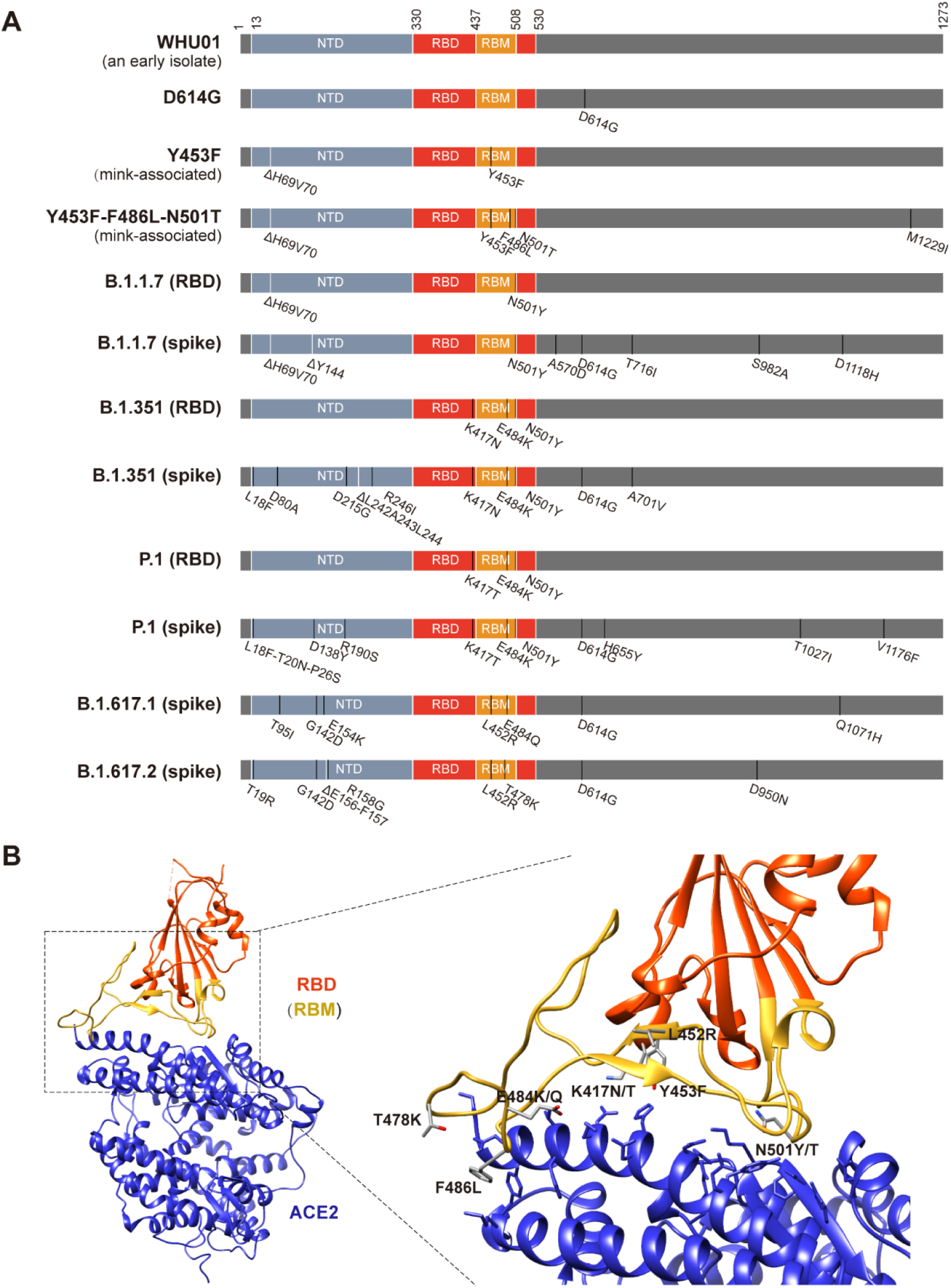
SARS-CoV-2 spike variants and related mutations investigated in the following studies. (**A**) Twelve SARS-CoV-2 spike genes investigated in this study are illustrated. The receptor binding domain (RBD) is in red and the receptor binding motif (RBM) is in yellow. Spike mutations associated with each variant are indicated. (**B**) Interactions between SARS-CoV-2 RBD (red) and ACE2 (blue) involve a large number of contact residues (PDB accession no. 6M0J). The so-called receptor binding motif (RBM) within the RBD is indicated in yellow. The ACE2 residues in less than 4 Å from RBD atoms are shown. All SARS-CoV-2 variant-associated RBD mutations investigated in this study are shown and labelled.

### Pseudoviruses of the B.1.1.7, B.1.351, and P.1 variants displayed patterns of animal-ACE2 tropism distinct from the patterns of other variants

To evaluate ACE2 ortholog-mediated viral entry of the above-mentioned SARS-CoV-2 variants, we produced luciferase reporter retroviruses pseudotyped with one of these different spike variants. These reporter pseudoviruses were used to infect 293T cells expressing each of eight ACE2 orthologs (**Figure 2 and Table 1**), including the human ACE2 and ACE2 orthologs of cattle (*Bos taurus*), pig (*Sus scrofa domesticus*), cat (*Felis catus*), rabbit (*Oryctolagus cuniculus*), bat (*Rhinolophus sinicus* isolate Rs-3357), rat (*Rattus norvegicus*), and mouse (*Mus musculus*). Parallel infection experiments using 293T cells transfected with an empty vector plasmid were included as controls. Consistent with our previous report^14^, the early isolate WHU01 efficiently infected 293T cells expressing most of the tested ACE2 orthologs except for that of rat and mouse (**Figure 2A**). While a similar pattern was also observed with the variants D614G, and Y453F (**Figure 2B and C**), distinct patterns were observed with the remaining variants (**Figure 2D-L**). Significant changes were observed with the VOCs B.1.1.7, B.1.351, and P.1, all of which showed ability to use ACE2 orthologs of rat and mouse for entry (**Figure 2E-J**). In addition, the variants B.1.351 and P.1 showed complete loss of the ability to use bat ACE2 for entry (**Figure 2G-J**). These data suggest that the mutations acquired by different variants might have changed SARS-CoV-2 RBD affinity to different ACE2 orthologs.

**Figure 2.**
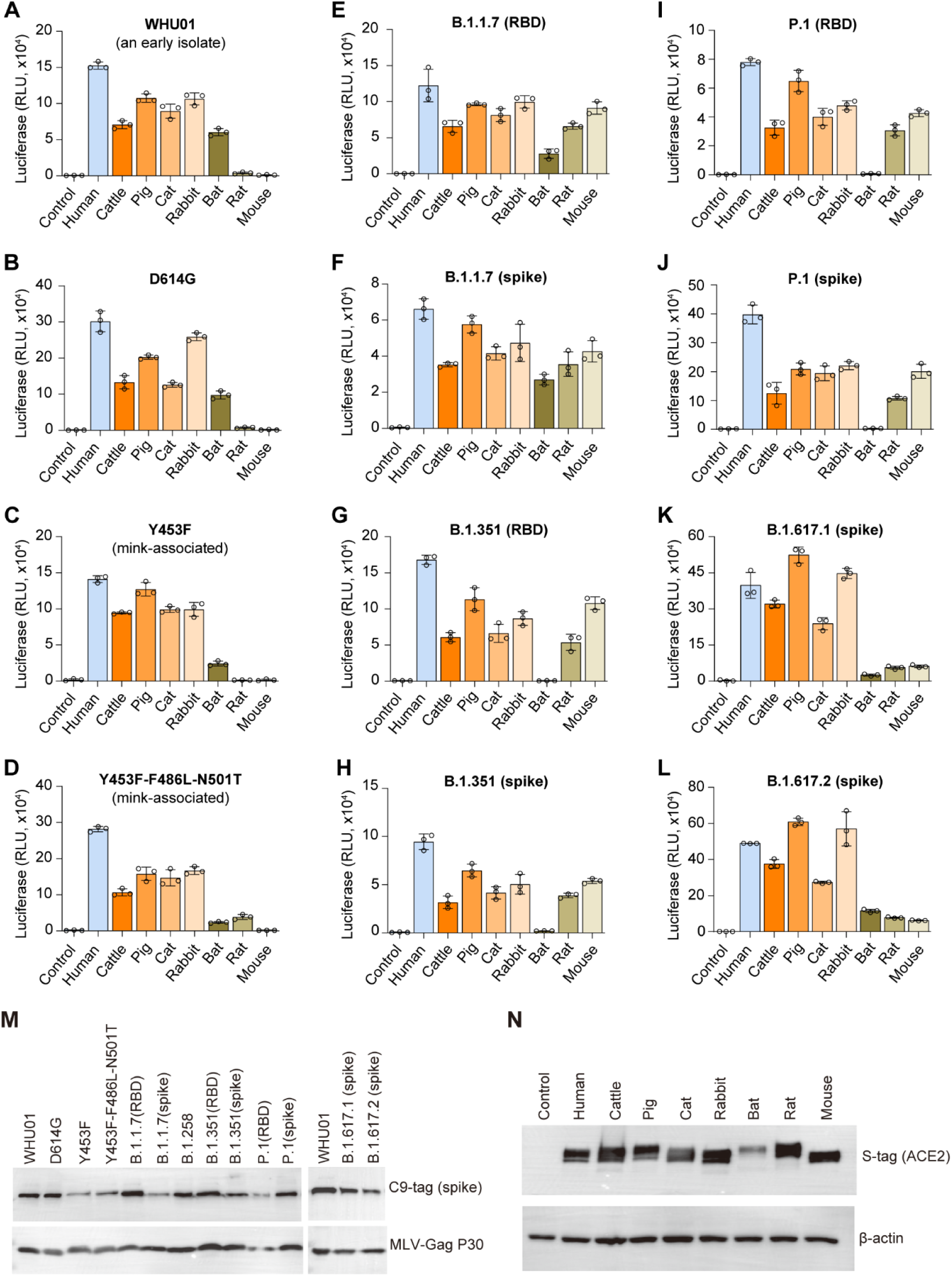
Animal-ACE2 tropism of SARS-CoV-2-variant pseudoviruses. **(A-L)** 293T cells expressing one of the eight indicated ACE2 orthologs were infected with each of twelve pseudoviruses corresponding to the twelve SARS-CoV-2 spike variants shown in Figure 1. ACE2-mediated pseudovirus entry was measured by a luciferase reporter expression at 48 hours post infection. Data shown are representative of three independent experiments performed by two different people with similar results, and data points represent mean ± s.d. of three biological replicates. (**M**) MLV retroviral vector-based pseudoviruses of the indicated SARS-CoV-2 variants were validated for the presence of spike protein (C9-tag) and MLV-Gag p30 protein. (**N**) 293T cells transfected with the indicated ACE2 genes (S-tagged) were collected at 48 hours post transfection. Expression levels of the indicated ACE2 proteins were detected using Western Blot.

**Table 1.**
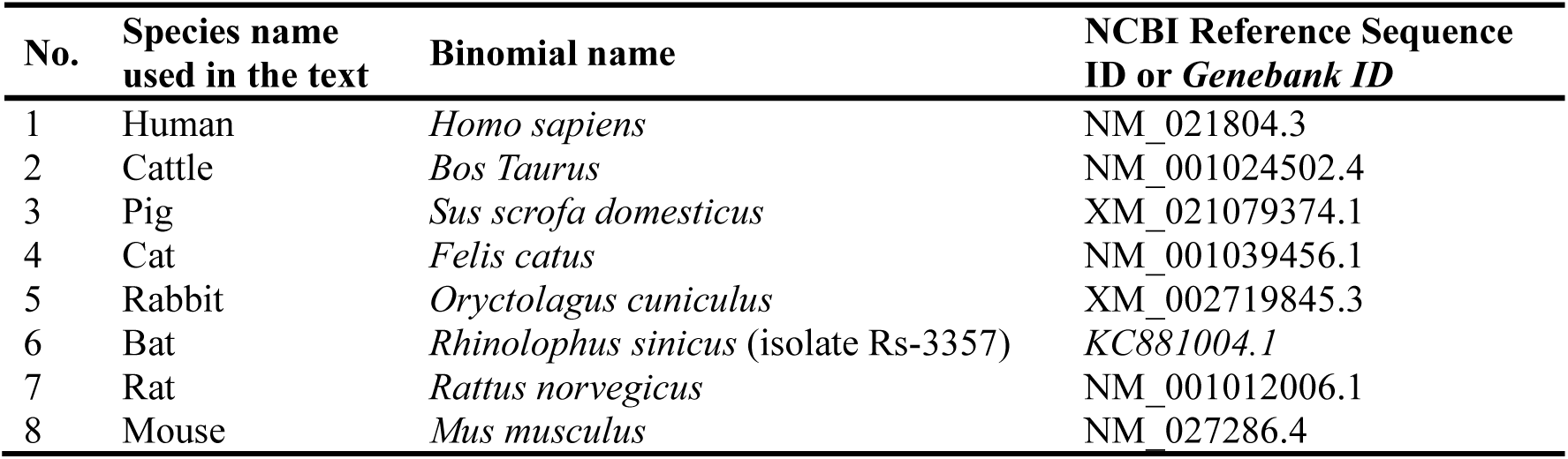
Species and accession numbers of the amino-acid sequences of the eight ACE2 orthologs investigated in this study.

### Y453F-RBD and the B.1.617.2-RBD have enhanced affinity to human ACE2

We then sought to quantitatively study the ability of the variants to utilize various animal orthologs of ACE2. The extracellular domains of the eight ACE2 orthologs and the RBD domains of seven spike variants, including WHU01, Y453F, B.1.1.7, B.1.351, P.1, B.1.617.1, and B.1.617.2, were expressed in 293F cells as immunoglobulin Fc fusion proteins. Purified ACE2 and RBD recombinant proteins were then used as immobilized receptors and analytes, respectively, in a bio-layer interferometry (BLI)-based assay to measure the interaction kinetics of fifty-six RBD-ACE2 pairs (**Figure 3 and Table 2**). When the RBD of early SARS-CoV-2 isolate WHU01 was tested against different ACE2 proteins, strong interactions were observed for human, cattle, pig, cat, and rabbit ACE2 proteins, with human ACE2 showing the highest affinity to this RBD. No any binding signal was detected for mouse ACE2 (**Figure 3H)**. We then compared the kinetics data of each RBD variants to the data of the WHU01 variant. We observed that the Y453F mutation increased RBD affinity to human ACE2 by ∼4.7-fold (**Figure 3A and Table 2**), consistent with a very recent report^42^. In addition, the B.1.617.2 RBD but not the B.1.617.1 RBD was also found with >2-fold higher affinity to human ACE2 (**Figure 3A and Table 2**), consistent with the fast spreading of this variant across the world.

**Figure 3.**
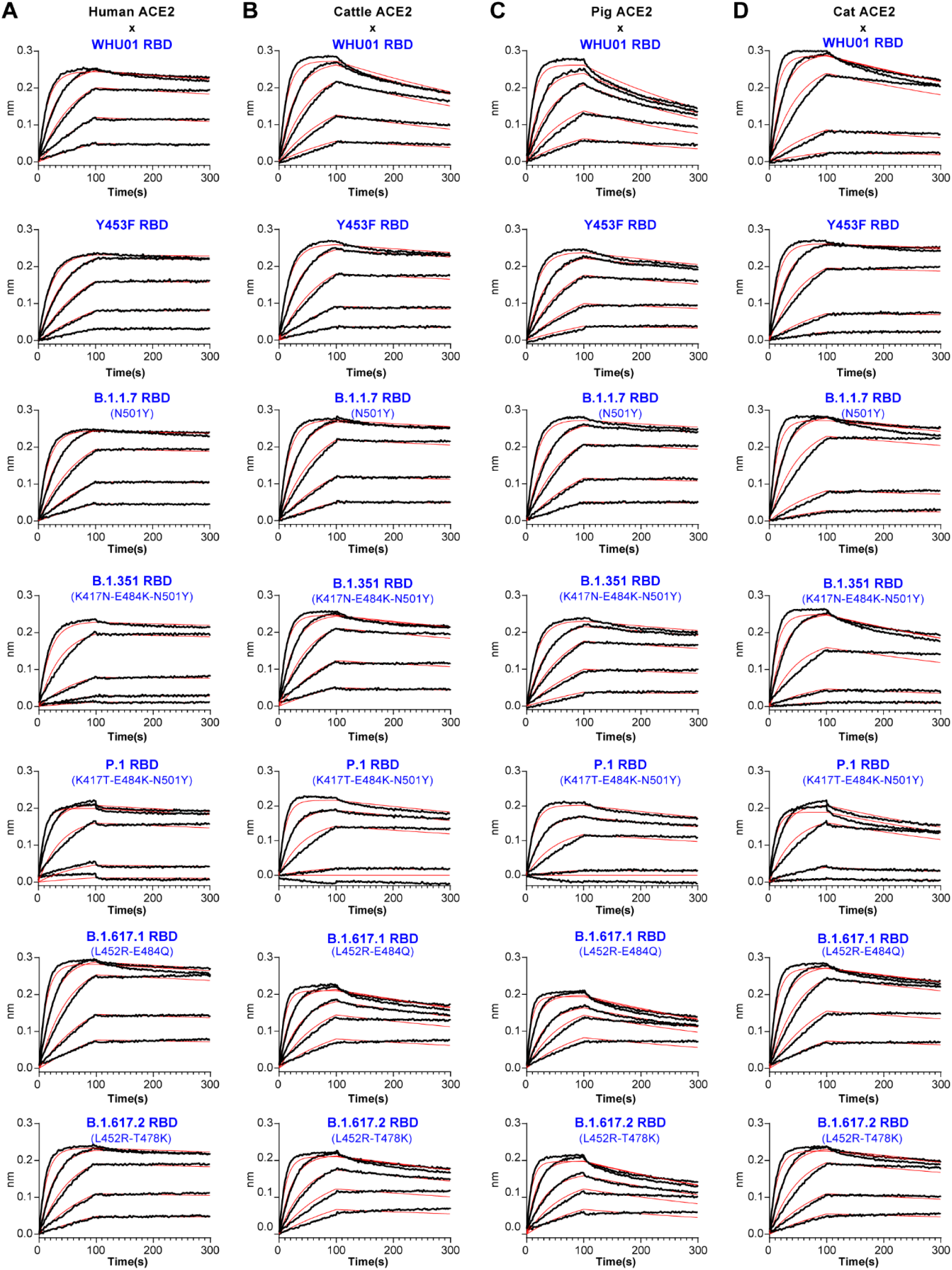
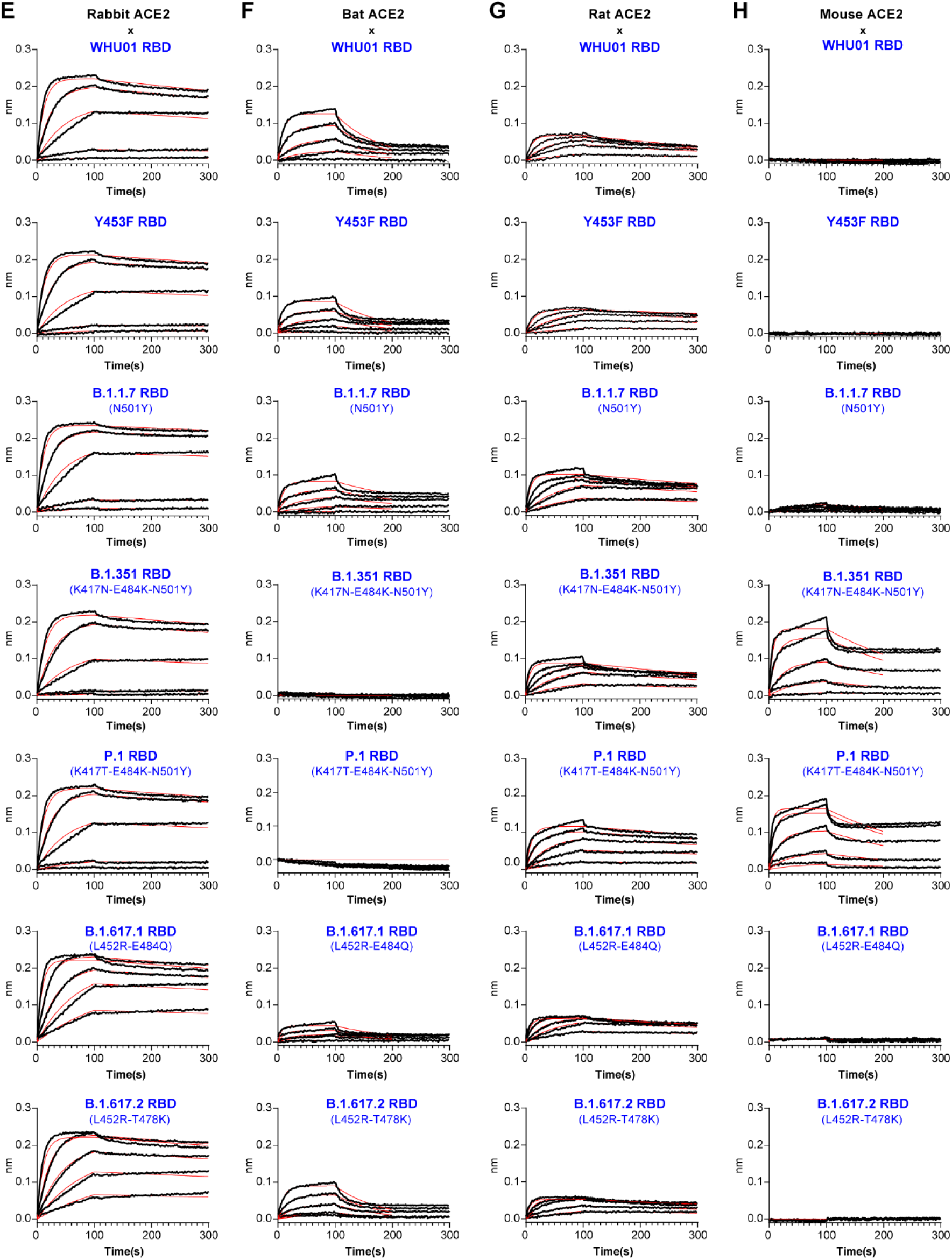
Interaction kinetics of fifty-six RBD-ACE2 pairs. (**A-H**) A bio-layer interferometry (BLI)-based assay was used to measure the kinetics of RBD-ACE2 interactions. Recombinant RBD proteins of seven SARS-CoV-2 variants, including WHU01, B.1.298, B.1.1.7, B.1.351, P.1, B.1.617.1, and B.1.617.2 were sequentially used as analytes to measure their interactions with recombinant ACE2 proteins of eight species, including human (A), cattle (B), pig (C), cat (D), rabbit (E), bat (F), rat (G), mouse (H). The black lines are the binding sensorgrams for each analyte at 100 nM, 50 nM, 25 nM, 12.5 nM, or 6.25 nM. The red lines show fits of the data to a 1:1 Langmuir binding model (global fit). Full kinetics data, including the on-rate (k_a_, M^-1^s^-1^), off-rate (k_dis_, s^-1^), and affinity (K_D_, nM) for all the fifty-six RBD-ACE2 interaction pairs, are provided in Table 2.

**Table 2.**
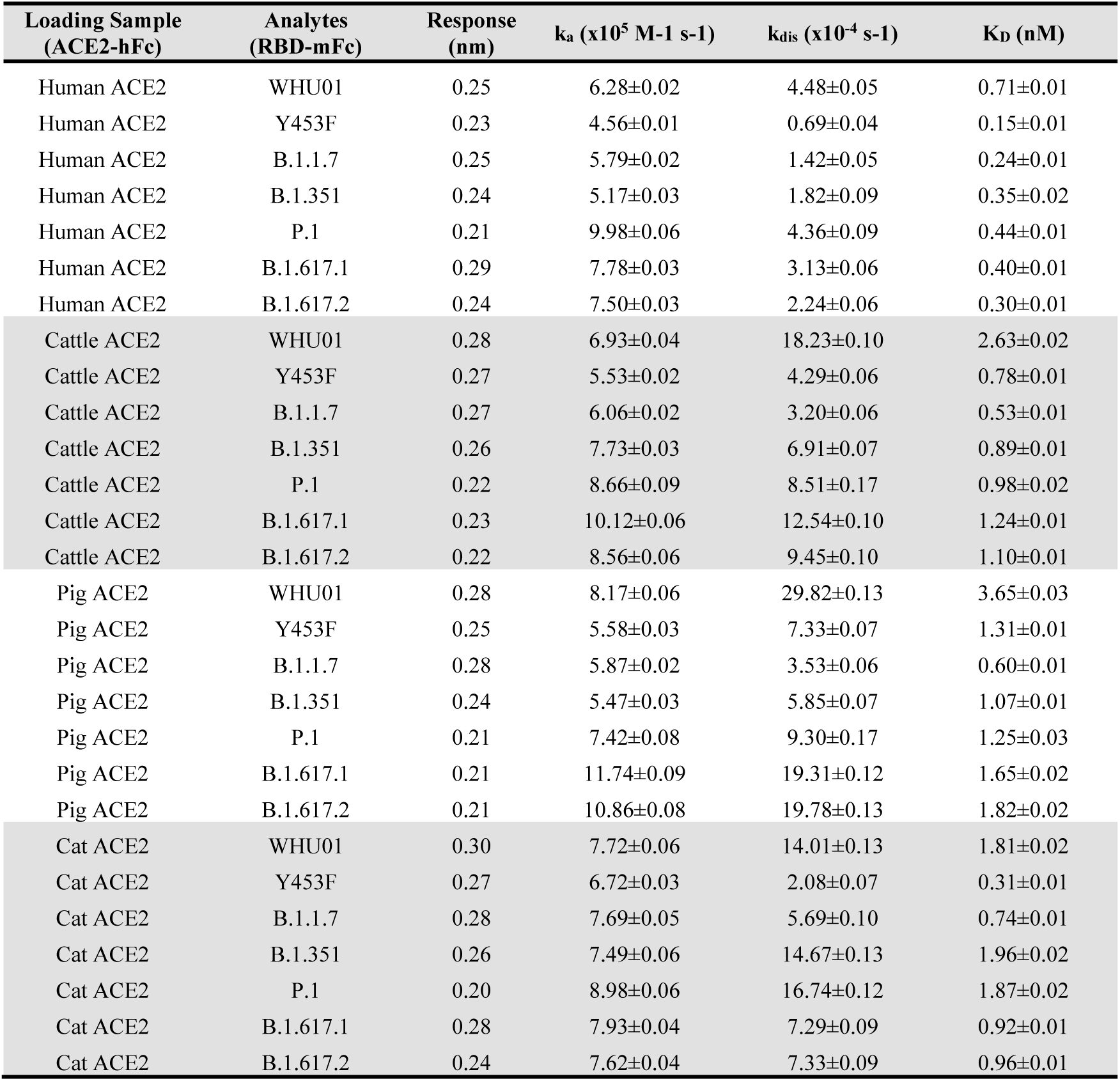
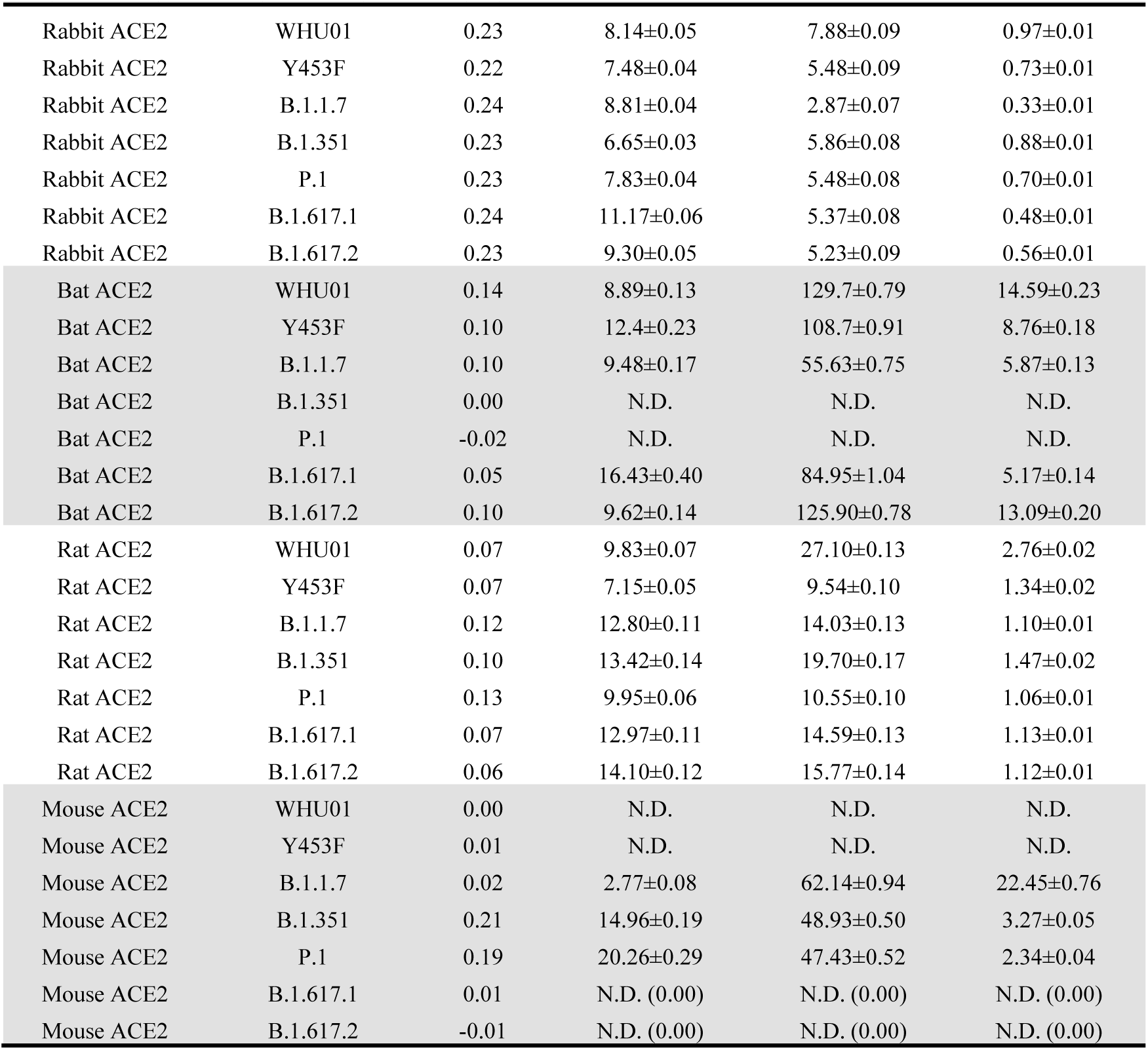
Interaction kinetics data, including the BLI Response (nm) at 100 nM analyte, on-rate (k_a_, M^-1^s^-1^), off-rate (k_dis_, s^-1^), and affinity (K_D_, nM), of the fifty-six ACE2-RBD pairs

| Loading Sample (ACE2-hFc) | Analytes (RBD-mFc) | Response (nm) | $k_a$ ($\times 10^5 M^{-1} s^{-1}$) | $k_{dis}$ ($\times 10^{-4} s^{-1}$) | $K_D$ (nM) |
| --- | --- | --- | --- | --- | --- |
| Human ACE2 | WHU01 | 0.25 | 6.28±0.02 | 4.48±0.05 | 0.71±0.01 |
| Human ACE2 | Y453F | 0.23 | 4.56±0.01 | 0.69±0.04 | 0.15±0.01 |
| Human ACE2 | B.1.1.7 | 0.25 | 5.79±0.02 | 1.42±0.05 | 0.24±0.01 |
| Human ACE2 | B.1.351 | 0.24 | 5.17±0.03 | 1.82±0.09 | 0.35±0.02 |
| Human ACE2 | P.1 | 0.21 | 9.98±0.06 | 4.36±0.09 | 0.44±0.01 |
| Human ACE2 | B.1.617.1 | 0.29 | 7.78±0.03 | 3.13±0.06 | 0.40±0.01 |
| Human ACE2 | B.1.617.2 | 0.24 | 7.50±0.03 | 2.24±0.06 | 0.30±0.01 |
| Cattle ACE2 | WHU01 | 0.28 | 6.93±0.04 | 18.23±0.10 | 2.63±0.02 |
| Cattle ACE2 | Y453F | 0.27 | 5.53±0.02 | 4.29±0.06 | 0.78±0.01 |
| Cattle ACE2 | B.1.1.7 | 0.27 | 6.06±0.02 | 3.20±0.06 | 0.53±0.01 |
| Cattle ACE2 | B.1.351 | 0.26 | 7.73±0.03 | 6.91±0.07 | 0.89±0.01 |
| Cattle ACE2 | P.1 | 0.22 | 8.66±0.09 | 8.51±0.17 | 0.98±0.02 |
| Cattle ACE2 | B.1.617.1 | 0.23 | 10.12±0.06 | 12.54±0.10 | 1.24±0.01 |
| Cattle ACE2 | B.1.617.2 | 0.22 | 8.56±0.06 | 9.45±0.10 | 1.10±0.01 |
| Pig ACE2 | WHU01 | 0.28 | 8.17±0.06 | 29.82±0.13 | 3.65±0.03 |
| Pig ACE2 | Y453F | 0.25 | 5.58±0.03 | 7.33±0.07 | 1.31±0.01 |
| Pig ACE2 | B.1.1.7 | 0.28 | 5.87±0.02 | 3.53±0.06 | 0.60±0.01 |
| Pig ACE2 | B.1.351 | 0.24 | 5.47±0.03 | 5.85±0.07 | 1.07±0.01 |
| Pig ACE2 | P.1 | 0.21 | 7.42±0.08 | 9.30±0.17 | 1.25±0.03 |
| Pig ACE2 | B.1.617.1 | 0.21 | 11.74±0.09 | 19.31±0.12 | 1.65±0.02 |
| Pig ACE2 | B.1.617.2 | 0.21 | 10.86±0.08 | 19.78±0.13 | 1.82±0.02 |
| Cat ACE2 | WHU01 | 0.30 | 7.72±0.06 | 14.01±0.13 | 1.81±0.02 |
| Cat ACE2 | Y453F | 0.27 | 6.72±0.03 | 2.08±0.07 | 0.31±0.01 |
| Cat ACE2 | B.1.1.7 | 0.28 | 7.69±0.05 | 5.69±0.10 | 0.74±0.01 |
| Cat ACE2 | B.1.351 | 0.26 | 7.49±0.06 | 14.67±0.13 | 1.96±0.02 |
| Cat ACE2 | P.1 | 0.20 | 8.98±0.06 | 16.74±0.12 | 1.87±0.02 |
| Cat ACE2 | B.1.617.1 | 0.28 | 7.93±0.04 | 7.29±0.09 | 0.92±0.01 |
| Cat ACE2 | B.1.617.2 | 0.24 | 7.62±0.04 | 7.33±0.09 | 0.96±0.01 |

**Table 2-continued.** Interaction kinetics data, including the BLI Response (nm) at 100 nM analyte, on-rate ( $k_a$ , $M^{-1}s^{-1}$ ), off-rate ( $k_{dis}$ , $s^{-1}$ ), and affinity ( $K_D$ , nM), of the fifty-six ACE2-RBD pairs
| Loading Sample<br>(ACE2-hFc) | Analytes<br>(RBD-mFc) | Response<br>(nm) | $k_a$ ($\times 10^5 M^{-1} s^{-1}$) | $k_{dis}$ ($\times 10^{-4} s^{-1}$) | $K_D$ (nM) |
| --- | --- | --- | --- | --- | --- |
| Rabbit ACE2 | WHU01 | 0.23 | 8.14±0.05 | 7.88±0.09 | 0.97±0.01 |
| Rabbit ACE2 | Y453F | 0.22 | 7.48±0.04 | 5.48±0.09 | 0.73±0.01 |
| Rabbit ACE2 | B.1.1.7 | 0.24 | 8.81±0.04 | 2.87±0.07 | 0.33±0.01 |
| Rabbit ACE2 | B.1.351 | 0.23 | 6.65±0.03 | 5.86±0.08 | 0.88±0.01 |
| Rabbit ACE2 | P.1 | 0.23 | 7.83±0.04 | 5.48±0.08 | 0.70±0.01 |
| Rabbit ACE2 | B.1.617.1 | 0.24 | 11.17±0.06 | 5.37±0.08 | 0.48±0.01 |
| Rabbit ACE2 | B.1.617.2 | 0.23 | 9.30±0.05 | 5.23±0.09 | 0.56±0.01 |
| Bat ACE2 | WHU01 | 0.14 | 8.89±0.13 | 129.7±0.79 | 14.59±0.23 |
| Bat ACE2 | Y453F | 0.10 | 12.4±0.23 | 108.7±0.91 | 8.76±0.18 |
| Bat ACE2 | B.1.1.7 | 0.10 | 9.48±0.17 | 55.63±0.75 | 5.87±0.13 |
| Bat ACE2 | B.1.351 | 0.00 | N.D. | N.D. | N.D. |
| Bat ACE2 | P.1 | -0.02 | N.D. | N.D. | N.D. |
| Bat ACE2 | B.1.617.1 | 0.05 | 16.43±0.40 | 84.95±1.04 | 5.17±0.14 |
| Bat ACE2 | B.1.617.2 | 0.10 | 9.62±0.14 | 125.90±0.78 | 13.09±0.20 |
| Rat ACE2 | WHU01 | 0.07 | 9.83±0.07 | 27.10±0.13 | 2.76±0.02 |
| Rat ACE2 | Y453F | 0.07 | 7.15±0.05 | 9.54±0.10 | 1.34±0.02 |
| Rat ACE2 | B.1.1.7 | 0.12 | 12.80±0.11 | 14.03±0.13 | 1.10±0.01 |
| Rat ACE2 | B.1.351 | 0.10 | 13.42±0.14 | 19.70±0.17 | 1.47±0.02 |
| Rat ACE2 | P.1 | 0.13 | 9.95±0.06 | 10.55±0.10 | 1.06±0.01 |
| Rat ACE2 | B.1.617.1 | 0.07 | 12.97±0.11 | 14.59±0.13 | 1.13±0.01 |
| Rat ACE2 | B.1.617.2 | 0.06 | 14.10±0.12 | 15.77±0.14 | 1.12±0.01 |
| Mouse ACE2 | WHU01 | 0.00 | N.D. | N.D. | N.D. |
| Mouse ACE2 | Y453F | 0.01 | N.D. | N.D. | N.D. |
| Mouse ACE2 | B.1.1.7 | 0.02 | 2.77±0.08 | 62.14±0.94 | 22.45±0.76 |
| Mouse ACE2 | B.1.351 | 0.21 | 14.96±0.19 | 48.93±0.50 | 3.27±0.05 |
| Mouse ACE2 | P.1 | 0.19 | 20.26±0.29 | 47.43±0.52 | 2.34±0.04 |
| Mouse ACE2 | B.1.617.1 | 0.01 | N.D. (0.00) | N.D. (0.00) | N.D. (0.00) |
| Mouse ACE2 | B.1.617.2 | -0.01 | N.D. (0.00) | N.D. (0.00) | N.D. (0.00) |

### N501Y mutation enhances RBD affinity to most of the tested ACE2 proteins, especially cattle and pig ACE2 proteins

In the case of the B.1.1.7 RBD which has a N501Y signature mutation, we observed 2.9-fold increase in the RBD affinity to human ACE2 (**Figure 3A and Table 2**). This is consistent with the binding data from other reports and the increased infectivity of the B.1.1.7 variant^39-41,43-45^. Structural studies show that the Tyr501 residue of the B.1.1.7 RBD inserts into a cavity at the binding interface and forms a perpendicular π–π stacking interaction with the Tyr41residue of ACE2^ref.45^. Perhaps because of a same mechanism, the B.1.1.7 RBD also displayed significantly increased affinity to all the other tested ACE2 proteins, except for bat ACE2 that has a His41 rather than a Tyr41 (**Figure 3B-H and Table 2**). More pronounced affinity increase was observed with cattle ACE2 (5.0-fold) and pig ACE2 (6.1-fold), making the affinities of cattle and pig ACE2 proteins to B.1.1.7 RBD slightly higher than that of human ACE2 to WHU01 RBD (**Figure 3B and C, and Table 2**). Only very weak binding signals were detected for mouse ACE2, though the B.1.1.7 pseudovirus efficiently utilized mouse ACE2 for entry (**Figure 2E, F, and 3H**), reflecting the sensitivity difference of the two assays for discriminating weak and strong interactions.

### The RBDs of B.1.351 and P.1 have increased affinities to cattle, pig, and mouse ACE2 proteins

In the case of the RBDs of B.1.351 and P.1 which share the E484K-N501Y signature mutations, almost identical patterns were observed (**Figure 3**). Compared to WHU01 RBD, B.1.351 and P.1 RBDs respectively also showed 2.9- and 2.7-fold higher affinities to cattle ACE2, and 3.4- and 2.9-fold higher affinities to pig ACE2 protein, possibly because of the shared N501Y mutation in these two variants (**Figure 3B and C, and Table 2**). On the other hand, consistent with the pseudovirus infection data (**Figure 2G-J**), the B.1.351 and P.1 RBDs showed a complete loss of interaction with bat ACE2, and a clear gain of interaction with mouse ACE2 (**Figure 3F and H**). Additional pseudovirus infection experiments showed that every single RBD mutation (K417N, K417T, E484K, or N501Y) of the B.1.351 and P.1 variants contribute to the utilization of mouse ACE2 as a functional receptor, with the N501Y mutation showed the most prominent effect (**Figure 4**). A Y41A single mutation in mouse ACE2 significantly impaired infectivity of all the single-spike-mutation pseudoviruses, but not the triple-spike-mutation pseudoviruses, while Y41A-H353A double mutation in mouse ACE2 significantly impaired infectivity of all the pseudoviruses, but much less pronounced effect was observed with the triple-spike-mutation pseudoviruses (**Figure 4**). These data suggest that the B.1.351 and P.1 RBDs gain binding to mouse ACE2 through the mutated RBD residues forming multiple new interactions with ACE2.

**Figure 4.**
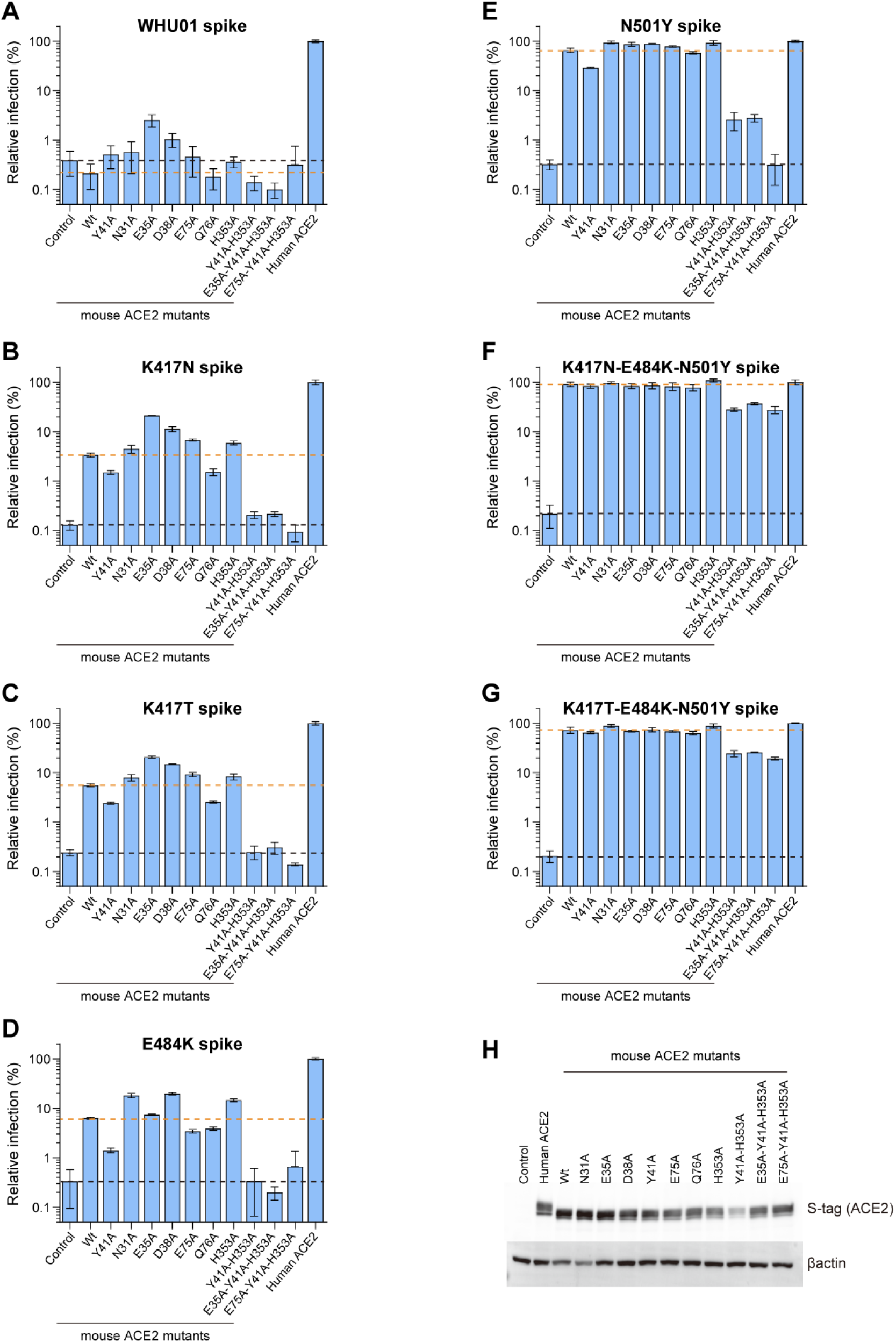
The effects of K417N, K417T, E484K, N501Y mutations on the ability of SARS-CoV-2 pseudovirus to utilize mouse ACE2. **(A-G)** SARS-CoV-2 pseudoviruses carrying the indicated spike mutations were used to infect 293T cells expressing human ACE2 or a mouse ACE2 mutant. 293T cells transfected with an empty vector were used as a control. ACE2-mediated pseudovirus entry was measured by a luciferase reporter expression at 48 hours post infection. Luminescence values were divided by the values observed with human ACE2 to calculate Relative Infection (%) values. (**H**) Expression levels of the ACE2 constructs used in the above experiments were detected using Western blotting. Data shown in A-G are representative of two independent experiments performed by two different people with similar results, and data points represent mean ± s.d. of three biological replicates.

### Spike mutations cause escape from potent neutralizing antibodies in clinical use

We then performed pseudovirus neutralization assays to evaluate sensitivity of these diverse SARS-CoV-2 variants to four therapeutic antibodies in clinical use (etesevimab/LY-CoV016, bamlanivimab/LY-CoV555, casirivimab/REGN10933, and imdevimab/REGN10987)^8-12^. While both the early strain WHU01 and the D614G variant were highly sensitive to all the four antibodies, all the RBD-mutated variants showed partial or complete resistance to at least one antibody (**Figure 5**). Specifically, both of the variants carrying a Y453F mutation showed strong resistance to REGN10933 (**Figure 5C and D**). The VOC B1.1.7 showed partial resistance to LY-CoV016 (**Figure 5E and F**). It is of note that the VOCs B.1.351 and P.1 showed strong resistance to REGN10933, and complete resistance to both LY-CoV016 and LY-CoV555, the two components of an antibody-cocktail therapy authorized for emergency use by the U.S. FDA^10-12^ (**Figure 5G-J**). The B.1.617.1 variant showed complete resistance to LY-CoV555 and partial resistance to REGN10933 (**Figure 5K)**. Although the VOC B.1.617.2 also showed strong resistance to LY-CoV555, it’s sensitivity to LY-CoV016 was found significantly increased (**Figure 5L**). Consistent with these neutralization data, analyzing structural data of these antibodies in complex with the original RBD revealed contact of LY-CoV016 with RBD residues Lys417, LY-CoV555 with RBD residues Glu484 and Phe486, REGN10933 with RBD residues Lys417, Tyr453, Glu484, and Phe486, and REGN10987 with RBD residues Asn439 and Gln498 (**Figure 5M-P**).

**Figure 5.**
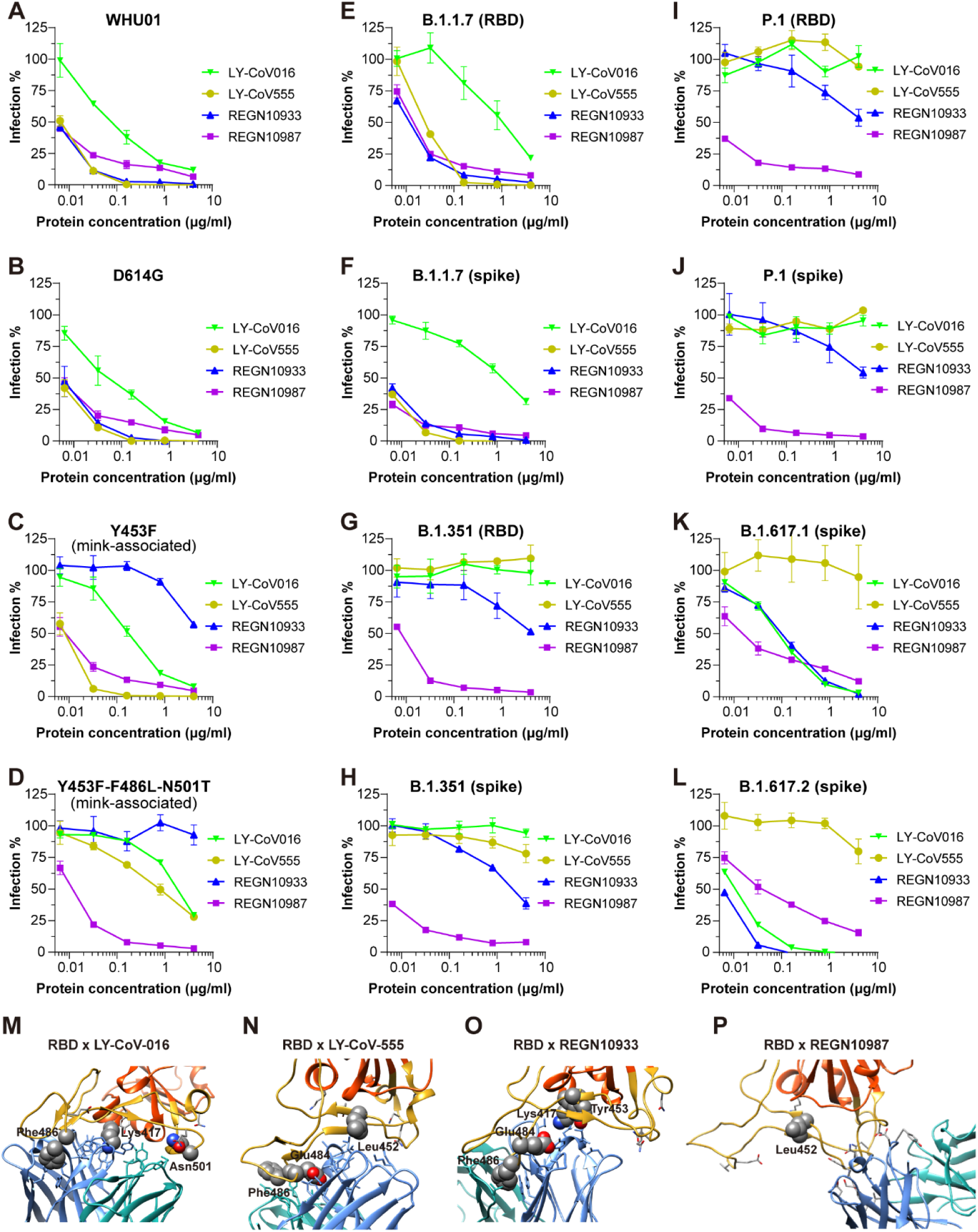
Neutralization sensitivity of SARS-CoV-2 variants to four monoclonal antibodies in clinical use. (**A-L**) Twelve SARS-CoV-2 variant pseudoviruses were tested for their neutralization sensitivity to four monoclonal antibodies (LY-CoV016/etesevimab, LY-CoV555/bamlanivimab, REGN10933/casirivimab, REGN10987/imdevimab) that constitute two antibody cocktails authorized by the U.S. FDA for emergency use. HeLa-hACE2 cells, a stable cell line that overexpresses human ACE2, were infected with each of the twelve pseudoviruses in the presence of the indicated neutralizing antibodies at the indicated concentrations. Viral entry was measured by luciferase reporter expression at 48 hours post infection. Luminescence values observed at each concentration were divided by the values observed at concentration zero to calculate percentage-of-infection (Infection%) values. Data shown are representative of three independent experiments performed by two different people with similar results, and data points represent mean ± s.d. of three biological replicates. (**M-P**) Structures of SARS-CoV-2 RBD in complex with LY-CoV016 (M; PDB accession no. 7C01), LY-CoV555 (N; PDB accession no. 7MKG), REGN10933 (O; PDB accession no. 6XDG), or REGN10987 (P; PDB accession no. 6XDG) are analyzed. The RBD and RBM are shown in red and yellow, respectively. Antibody heavy chain and light chain are shown in blue and green, respectively. Antibody residues in less than 4 Å from RBD atoms and RBD residues associated with SARS-CoV-2 mutations are shown as sticks. SARS-CoV-2 mutation-associated RBD residues that have direct contact with antibody residues or reside within 5 Å from antibody atoms are shown as spheres and labelled.

### The impact of RBD mutations on antibody affinity

We then performed BLI assays to quantitatively measure the kinetics of antibody-RBD interactions (**Figure 6 and Table 3**). The binding data are fully consistent with the pseudovirus neutralization data. Specifically, the mink-associated Y453F mutation decreased RBD affinity to REGN10933 (**Figure 6C**). The B.1.1.7-signature mutation N501Y decreased RBD binding to LY-CoV016 (**Figure 6A**). The B.1.351 RBD showed significantly decreased binding to REGN10933, and complete loss of interaction with LY-CoV016 and LY-CoV555 (**Figure 6A-C**). The P.1 RBD showed significantly decreased binding to REGN10933, weakly-detectable interaction with LY-CoV016, and complete loss of interaction with LY-CoV555 (**Figure 6A-C**). The B.1.617.1 RBD showed complete loss of interaction with LY-CoV555 (**Figure 6B**). Consistent with the neutralization data, the B.1.617.2 RBD showed significantly decreased affinity to LY-Cov555, but increased affinity to REGN10933 (**Figure 5L, 6B, and 6C**). These data demonstrate that SARS-CoV-2 variants could easily develop resistance to neutralization antibodies or even antibody cocktails in clinical use, highlighting the necessity of developing broad-spectrum anti-coronavirus agents.

**Figure 6.**
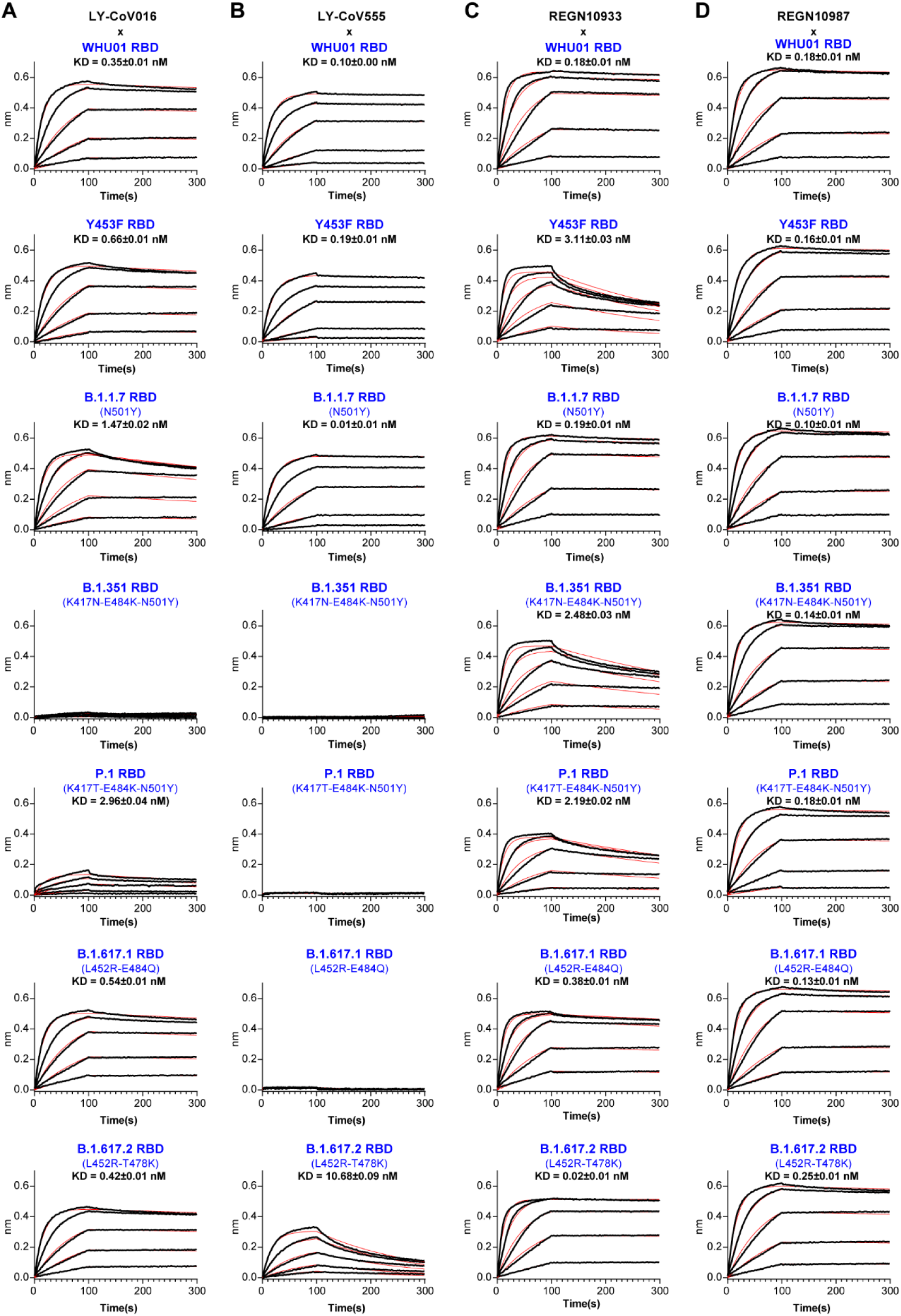
Interaction kinetics of twenty-eight RBD-antibody pairs. (**A-D**) The BLI-based assay was used to measure the kinetics of RBD-antibody interactions. Recombinant RBD proteins of seven SARS-CoV-2 spike variants, including WHU01, Y453F, B.1.1.7, B.1.351, P.1, B.1.617.1, and B.1.617.2 were sequentially used as analytes to measure their interactions with four monoclonal antibodies, including LY-CoV016 (A), LY-CoV555 (B), REGN10933 (C), and REGN10987 (D). The black lines are the binding sensorgrams for each analyte at 100 nM, 50 nM, 25 nM, 12.5 nM, or 6.25 nM. The red lines show fits of the data to a 1:1 Langmuir binding model (global fit). The affinity (K_D_, nM) for each interaction is indicated in the insets of the figures. Full kinetics data, including the on-rate (k_a_, M^-1^s^-1^), off-rate (k_dis_, s^-1^), and affinity (K_D_, nM) for each interaction, are provided in Table 3.

**Table 3.** Interaction kinetics data, including the BLI Response (nm) at 100 nM analyte, on-rate (k_a_, M^-1^s^-1^), off-rate (k_dis_, s^-1^), and affinity (K_D_, nM), of the twenty-eight mAb-RBD pairs

| Loading Sample (Antibody) | Analytes (RBD-mFc) | Response (nm) | $k_a$ ($\times 10^5 M^{-1} s^{-1}$) | $k_{dis}$ ($\times 10^{-4} s^{-1}$) | $K_D$ (nM) |
| --- | --- | --- | --- | --- | --- |
| LY-CoV016 | WHU01 | 0.57 | 5.98±0.01 | 2.10±0.04 | 0.35±0.01 |
| LY-CoV016 | Y453F | 0.51 | 5.51±0.02 | 3.62±0.05 | 0.66±0.01 |
| LY-CoV016 | B.1.1.7 | 0.52 | 6.31±0.03 | 9.25±0.08 | 1.47±0.02 |
| LY-CoV016 | B.1.351 | 0.03 | N.D. | N.D. | N.D. |
| LY-CoV016 | P.1 | 0.16 | 5.19±0.05 | 15.37±0.16 | 2.96±0.04 |
| LY-CoV016 | B.1.617.1 | 0.52 | 6.13±0.02 | 3.33±0.04 | 0.54±0.01 |
| LY-CoV016 | B.1.617.2 | 0.46 | 5.95±0.01 | 2.50±0.04 | 0.42±0.01 |
| LY-CoV555 | WHU01 | 0.50 | 6.30±0.01 | 0.62±0.03 | 0.10±0.00 |
| LY-CoV555 | Y453F | 0.45 | 5.53±0.01 | 1.06±0.03 | 0.19±0.01 |
| LY-CoV555 | B.1.1.7 | 0.49 | 5.12±0.01 | 0.06±0.03 | 0.01±0.01 |
| LY-CoV555 | B.1.351 | 0.00 | N.D. | N.D. | N.D. |
| LY-CoV555 | P.1 | 0.02 | N.D. | N.D. | N.D. |
| LY-CoV555 | B.1.617.1 | 0.02 | N.D. | N.D. | N.D. |
| LY-CoV555 | B.1.617.2 | 0.33 | 5.09±0.04 | 54.35±0.18 | 10.68±0.09 |
| REGN10933 | WHU01 | 0.63 | 7.26±0.03 | 1.27±0.06 | 0.18±0.01 |
| REGN10933 | Y453F | 0.50 | 9.92±0.09 | 30.82±0.17 | 3.11±0.03 |
| REGN10933 | B.1.1.7 | 0.62 | 7.81±0.02 | 1.52±0.05 | 0.19±0.01 |
| REGN10933 | B.1.351 | 0.50 | 9.03±0.07 | 22.4±0.14 | 2.48±0.03 |
| REGN10933 | P.1 | 0.40 | 8.47±0.06 | 18.52±0.12 | 2.19±0.02 |
| REGN10933 | B.1.617.1 | 0.51 | 9.07±0.03 | 3.45±0.06 | 0.38±0.01 |
| REGN10933 | B.1.617.2 | 0.52 | 8.51±0.03 | 0.14±0.05 | 0.02±0.01 |
| REGN10987 | WHU01 | 0.66 | 6.09±0.02 | 1.10±0.04 | 0.18±0.01 |
| REGN10987 | Y453F | 0.62 | 5.49±0.01 | 0.90±0.03 | 0.16±0.01 |
| REGN10987 | B.1.1.7 | 0.66 | 6.12±0.01 | 0.60±0.04 | 0.10±0.01 |
| REGN10987 | B.1.351 | 0.64 | 6.42±0.02 | 0.90±0.04 | 0.14±0.01 |
| REGN10987 | P.1 | 0.57 | 5.90±0.02 | 1.09±0.05 | 0.18±0.01 |
| REGN10987 | B.1.617.1 | 0.67 | 6.41±0.02 | 0.80±0.04 | 0.13±0.01 |
| REGN10987 | B.1.617.2 | 0.62 | 5.96±0.02 | 1.51±0.04 | 0.25±0.01 |

### Spike mutations change sensitivity of SARS-CoV-2 variants to different ACE2-Ig constructs

ACE2-Ig is a recombinant Fc fusion protein of soluble human ACE2. Recently we developed a panel of ACE2-Ig constructs that potently neutralize SARS-CoV-2 early isolate and three related but distinct coronaviruses, demonstrating that ACE2-Ig is a promising broadly anti-coronavirus drug candidate ^14^. Here we evaluated *in vitro* neutralization sensitivity of the twelve SARS-CoV-2 variants to eight representative ACE2-Ig constructs previously described in three different studies, including three constructs from our recent study (ACE2-Ig-v1, ACE2-Ig-v1.1, ACE2-Ig-v3)^14^, one construct from Chan *et al* (ACE2-Ig-Chan-v2.4)^46^, and four constructs from Glasgow *et al* (ACE2-Ig-Glasgow-293, ACE2-Ig-Glasgow-310, ACE2-Ig-Glasgow-311, ACE2-Ig-Glasgow-313)^47^. Because we previously found that the CLD domain of human ACE2 has an ∼20-fold contribution to ACE2-Ig’s neutralization activity against SARS-CoV-2 pseudoviruses^14^, we used CLD-containing soluble ACE2 domains for all the constructs tested in this study. We got three interesting findings here (**Figure 7**). First, all the twelve SARS-CoV-2 variants were potently neutralized by all the eight ACE2-Ig constructs, demonstrating that ACE2-Ig is a broad-spectrum anti-SARS-CoV-2 agent. Second, the mink-associated variant Y453F-F486L-N501T showed partial resistance to ACE2-Ig-Glasgow-310 and ACE2-Ig-Glasgow-313, but clearly increased sensitivity to ACE2-Ig-v1, ACE2-Ig-v1.1 and ACE2-Ig-v3, the constructs that are not heavily mutated in the spike-binding interface of the soluble ACE2 domain (**Figure 7A-C, F, H**). These data suggest that extensively mutating the ACE2 residues near the RBD-binding interface should be avoided. It might result in compromised neutralization breadth, not to mention the risk of eliciting anti-drug antibody (ADA) immune response. Third, compared to the early isolate WHU01, most circulating variants, including the four VOCs B.1.1.7, B.1.351, P.1, and B.1.617.2 showed significantly increased (up to ∼15-fold) neutralization sensitivity to ACE2-Ig-v1, ACE2-Ig-v1.1 and ACE2-Ig-v3, the constructs that carry a wild-type or a D30E-mutated ACE2 domain (**Figure 7A-C**).

**Figure 7.**
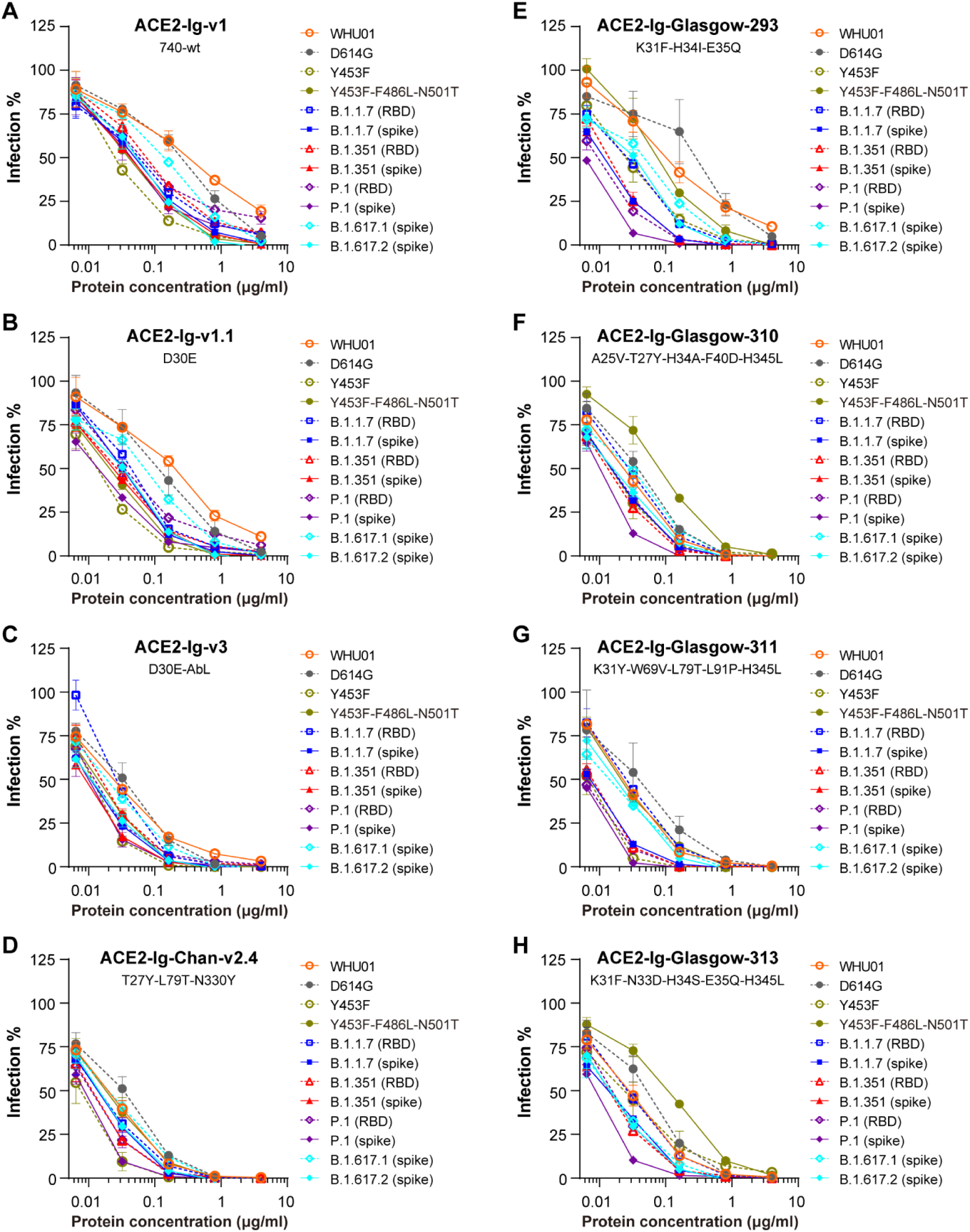
Neutralization sensitivity of SARS-CoV-2 variants to eight ACE2-Ig constructs. (**A-H**) Eight recombinant ACE2-Ig protein constructs, including three from our recent study (ACE2-Ig-v1, ACE2-Ig-v1.1, ACE2-Ig-v3; A-C) ^ref.14^, one from Chan *et al* (ACE2-Ig-Chan-v2.4; D) ^ref.46^; and four from Glasgow *et al* (ACE2-Ig-Glasgow-293, ACE2-Ig-Glasgow-310, ACE2-Ig-Glasgow-311, ACE2-Ig-Glasgow-313; E-H) ^ref.47^, were tested for their neutralization potencies against the twelve SARS-CoV-2 variant pseudoviruses. HeLa-hACE2 cells were infected with each of the twelve pseudoviruses in the presence of the indicated ACE2-Ig proteins at the indicated concentrations. Viral entry was measured by luciferase reporter expression at 48 hours post infection. Luminescence values observed at each concentration were divided by the values observed at concentration zero to calculate percentage-of-infection (Infection%) values. Data shown are representative of three independent experiments performed by two different people with similar results, and data points represent mean ± s.d. of three biological replicates.

## Discussion

In this study, we parallelly investigated multiple circulating SARS-CoV-2 variants, including two mink-associated variants, the B.1.258 and B.1.617 variants, and the VOCs B.1.1.7, B.1.351, and P.1. As receptor usage is a critical determinant for the host range of coronaviruses, we first investigated the ability of these variants to utilize animal orthologs of ACE2 as an entry receptor. We found that, in contrast to an early isolate WHU01, the VOCs B.1.351, and P.1 were able to bind and use mouse ACE2 for entry, suggesting that these variants may have gained ability to infect mouse (**Figure 2G-J and 3H**). Consistent with our finding, during the preparation of this manuscript, an *in vivo* study reported that the VOCs B.1.351 and P.1 were able to infect mouse and replicate to high titers in the lungs^48^. Mice are vaccine-inaccessible rodent species that have large population size and could access the territories of both humans and domestic animals. The fact that the B.1.351 and P.1 variants have gained ability to infect mice raises the possibility of wild rodents becoming a second SARS-CoV-2 reservoir, adding one more concerning factor to these variants.

In addition, we found that multiple variants, especially the VOCs B.1.1.7, B.1.351, and P.1, have significantly increased affinity to cattle and pig ACE2 proteins (**Figure 3 and Table 1**). Multiple previous studies have shown that cattle and pigs are not or only weakly susceptible to experimental infection with SARS-CoV-2^15,49-52^. Consistently, here we found that both cattle and pig ACE2 proteins have significantly lower affinities than that of human ACE2 to the RBD of an early isolate WHU01. However, in the case of the RBDs of the three VOCs, binding affinities of cattle and pig ACE2 proteins to these RBDs are comparable to that of human ACE2 to WHU01 RBD (**Table 2**). Cattle and pig ACE2 affinities to B.1.1.7 RBD are even slightly higher than that of human ACE2 to WHU01 RBD. It has been recently reported that the RBDs in the B.1.1.7 and B.1.351 spike proteins are more accessible than that of the early isolate^43,44^. This could further enable these variants to efficiently use cattle and pig ACE2 proteins. These data altogether suggest that cattle and pigs might now become susceptible to SARS-CoV-2. Cattle and pigs are extremely important livestock animals which serve as two major sources of meat for humans. It might be necessary to perform *in vivo* studies to re-evaluate the susceptibility of these species to SARS-CoV-2 variants, especially the widely circulating VOCs. On the other hand, the SARS-CoV-2 infected population is already in a huge size, with over 160 million confirmed infections worldwide. As of 29 March 2021, the VOC B.1.1.7 comprises roughly 95% of new SARS-CoV-2 infections in England. It has now been identified in at least 114 countries and exhibits a similar transmission increase (59 to 74%) in Denmark, Switzerland, and the United States^39-41^. So far, the VOCs B.1.351 and P.1 have also been identified in at least 68 and 37 countries, respectively. The enormous number of SARS-CoV-2 infection cases and high prevalence of these VOCs worldwide make it more and more necessary to closely monitor cattle and pigs for SARS-CoV-2 natural infections.

Our data about neutralization sensitivity of the SARS-CoV-2 variants to four monoclonal antibodies are consistent with multiple other studies^53-58^. Although, compared to the early isolate WHU01, all the tested variants carrying mutations in the RBD region showed partial or fully escape from at least one and up to three of the four tested antibodies (**Figure 5 and 6**), none of the variants escaped ACE2-Ig (**Figure 7**), a broad-spectrum anti-SARS-CoV-2 drug candidate. More importantly, we found that most of the tested SARS-CoV-2 variants had increased (up to ∼15-fold) sensitivity to ACE2-Ig constructs that carry a wild-type or a single-point-mutated ACE2 domain (**Figure 7A-B**). This might be explained by increased affinity of SARS-CoV-2 variant to human ACE2 or increased accessibility of the RBDs of SARS-CoV-2 variants. Indeed, we observed a 4.7-fold, 2.9-fold, 2.0-fold and 1.6-fold increase in the affinity of human ACE2 to the RBDs of the mink-associated Y453F variant, the B.1.1.7 variant, the B.1.351 variant, and the P.1 variant, respectively (**Table 2**). On the other hand, recent structural studies have shown that spike trimers of multiple SARS-CoV-2 variants, including a mink-associate variant and the VOCs B.1.1.7, B.1.1.28, and B1.351, have significantly higher propensity to adopt ‘RBD-up’ or open state than the D614G variant does^43,44^. These data suggest that SARS-CoV-2 was evolving toward better utilization of ACE2 as a receptor, either through increasing RBD affinity to ACE2, or through better exposure of its RBDs, or through both. It further indicates that ACE2 is still likely an essential receptor for SARS-CoV-2. ACE2 also serves as a cellular receptor for a number of other coronaviruses, including SARS-CoV, HCoV-NL63, Pangolin-CoV-2020, Bat-CoV RaTG13, and some other SARS-like CoVs found in bats^14,16,59^. ACE2-Ig is therefore a promising broadly anti-coronavirus drug candidate that might be used to treat and prevent infection of these diverse coronaviruses and their variants. It might also be a good alternative anti-SARS-CoV-2 agent for the populations who are not responsive to, or don’t have access to any prophylactic vaccines.

There are a couple of limitations of the current study. SARS-CoV-2 spike-pseudotyped reporter viruses, instead of live viruses, were used to study virus-ACE2 interaction and neutralization sensitivity of multiple circulating SARS-CoV-2 variants. It is difficult to obtain or access live SARS-CoV-2 variants, and even more difficult to simultaneously obtain multiple VOC live viruses and the mink-associated variant. Utilization of pseudotyped reporter virus to study virus-receptor interaction and neutralization is a well-established and widely used method in the field for studying various enveloped viruses, such as HIV^60,61^, influenza virus^62-64^, and coronaviruses including SARS-CoV-2^ref.14,34,47,64-68^. Findings about virus-receptor interaction and neutralization based on pseudovirus systems normally correlate well with that based on live viruses. In addition, we applied quantitative binding kinetics assays to validate all the key findings based on pseudovirus infection assays. This largely compensate the limitation of using pseudoviruses. On the other hand, ACE2-Ig is a promising broadly anti-SARS-CoV-2 drug candidate. More studies under *in vivo* conditions will be necessary to validate its potential for clinical applications. We have recently obtained a preliminary result showing that a single-dose injection or intranasal administration of an ACE2-Ig construct could significantly lower the viral load in the lungs of a COVID-19 mouse model (unpublished data). More mouse studies to test and optimize the protein’s *in vivo* pharmacokinetic properties and anti-SARS-CoV-2 efficacy are ongoing.

## Materials and Methods

### Cells

293T cells and HeLa cells were kindly provided by Stem Cell Bank, Chinese Academy of Sciences, confirmed mycoplasma-free by the provider, and maintained in Dulbecco’s Modified Eagle Medium (DMEM, Life Technologies) at 37 °C in a 5% CO_2_-humidified incubator. Growth medium was supplemented with 2 mM Glutamax-I (Gibco, Cat. No. 35050061), 100 µM non-essential amino acids (Gibco, Cat. No. 11140050), 100 U/mL penicillin and 100 µg/mL streptomycin (Gibco, Cat. No. 15140122), and 10% heat-inactivated FBS (Gibco, Cat. No. 10099141C). HeLa-based stable cells expressing human ACE2 were maintained under the same culture condition as HeLa, except that 3 µg/mL of puromycin was added to the growth medium. 293F cells for recombinant protein production were generously provided by Dr. Yu J. Cao (School of Chemical Biology and Biotechnology, Peking University Shenzhen Graduate School) and maintained in SMM 293-TII serum-free medium (Sino Biological, Cat. No. M293TII) at 37 °C, 8% CO_2_, in a shaker incubator at 125 rpm.

### Plasmids

DNA fragment encoding spike protein of SARS-CoV-2 WHU01 (GenBank: MN988668.1) was synthesized by the Beijing Genomic Institute (BGI, China) and then cloned into pCAGGS plasmid between EcoRI and XhoI restriction sites. Plasmids encoding SARS-CoV-2 spike variants were generated according to the in-fusion cloning protocol. To facilitate SARS-CoV-2 pseudovirus production, spike sequences for WHU01 and all the variants investigated in this study all contain a furin-cleavage site mutation (ΔPRRA) or a GSAS substitution of the PRRA furin-cleavage site. We had shown in our previous study that the ΔPRRA mutation does not affect SARS-CoV-2 cross-species receptor usage or neutralization sensitivity^14^. The retroviral reporter plasmids encoding a Gaussia luciferase reporter gene were constructed by cloning the reporter genes into the pQCXIP plasmid (Clontech). DNA fragments encoding C-terminally S-tagged ACE2 orthologs were synthesized in pUC57 backbone plasmid by Sangon Biotech (Shanghai, China). These fragments were then cloned into pQCXIP plasmid (Clontech) between SbfI and NotI restriction sites. Plasmids encoding recombinant RBD and soluble ACE2 variants were generated by cloning each of the gene fragments into a pCAGGS-based mouse-IgG2a or human IgG1 Fc fusion protein expression plasmid between NotI and BspEI sites. DNA fragments encoding heavy and light chains of anti-SARS-CoV-2 antibodies were synthesized by Sangon Biotech (Shanghai, China) and then cloned into a pCAGGS plasmid. Four antibodies (LY-CoV016, LY-CoV-555, REGN10933, and REGN10987)^ref.8-12^ that constitute two antibody cocktails authorized by the U.S. FDA for emergency use were included in this study.

### Western Blot to detect S-tagged ACE2 (ACE2-S-tag) expression in 293T cells

293T cells at 30% density in each well of 96 well plates were reverse transfected with 0.60 µg of plasmid in complex with 0.15 µL of lipofectamine 2000 (Life Technologies, Cat. No. 11668019). Twenty-four hours after transfection, cells in each well were lysed with 40 µL lysis buffer and 5 µL of the lysate was used for western blot. ACE2-S-tag expression was detected by using a mouse anti-S-tag monoclonal antibody 6.2 (Invitrogen, Cat. No. MA1-981), and an HRP-conjugated goat anti-mouse IgG Fc secondary antibody (Invitrogen, Cat. No. 31437). Beta-actin was used as an internal control.

### Production of reporter retroviruses pseudotyped with SARS-CoV-2 spike variants

MLV retroviral vector-based SARS-CoV-2 spike pseudotypes were produced according to our previous study^14^, with minor changes. In brief, 293T cells were seeded at 30% density in 150 mm dish at 12-15 hours before transfection. Cells were then transfected with 67.5 µg of polyethylenimine (PEI) Max 40,000 (Polysciences, Inc, Cat. No. 24765-1) in complex with 3.15 µg of plasmid encoding a spike variant, 15.75 µg of plasmid encoding murine leukemia virus (MLV) Gag and Pol proteins, and 15.75 µg of a pQCXIP-based luciferase reporter plasmid. Eight hours after transfection, cell culture medium was refreshed and changed to growth medium containing 2% FBS (Gibco, Cat. No. 10099141C) and 25 mM HEPES (Gibco, Cat. No. 15630080). Cell culture supernatants were collected at 36-48 hours post transfection, spun down at 3000×g for 10 min, and filtered through 0.45 µm filter units to remove cell debris. SARS-CoV-2 spike-pseudotyped viruses were then concentrated 10 times at 2000×g using 100 kDa cut-off Amicon Ultra-15 Centrifugal Filter Units (Millipore. Cat. No. UFC910024).

### Pseudovirus Titration

Pseudovirus titer were determined by TCID_50_ followed a previous protocol^69^. Viruses were diluted 100 times as a working solution and then serially diluted in a ½ log10 manner. Human ACE2 expressed HeLa cells were infected with those diluted viruses in 96-well plates. Culture supernatants were refreshed every 12 hours and loaded to a Gaussia luciferase assay at 48 hours post infection. TCID_50_ of each pseudovirus was calculated by the Reed-Muench method.

### Western Blot to detect spike protein (C9-tag) and MLV P30 protein in SARS-CoV-2 pseudovirus-containing supernatant

Pseudovirus-containing supernatants (8000 TCID50 of each pseudovirus) were mixed with SDS-PAGE loading buffer and boiled at 95°C for 10min. Spike protein (C9-tag) on the surface of pseudovirus was detected using a mouse anti-C9-tag monoclonal antibody 1D4 (Invitrogen, Cat. No. MA1-722), and an HRP-conjugated goat anti-mouse IgG Fc secondary antibody (Invitrogen, Cat. No. 31437). MLV P30 protein within the pseudovirus capsid was detected using a rabbit anti-MLV-P30 polyclonal antibody (Origene, Cat. No. AP33447PU-N) and an HRP-conjugated goat anti-rabbit IgG Fc secondary antibody (Invitrogen, Cat. No. 31463).

### SARS-CoV-2 pseudovirus infection of 293T cells expressing ACE2 orthologs

Pseudovirus infection assay was performed according to our previous study^14^. In brief, 293T cells at 30% density in each well of gelatin (Millipore, ES-006-B) pre-coated 96-well plates were reverse transfected with 0.15 µL of lipofectamine 2000 (Life Technologies, Cat. No. 11668019) in complex with 60 ng of a vector control plasmid or a plasmid encoding ACE2 orthologs or ACE2 mutants. Twenty-four hours later, cells in each well were infected with 2000 TCID_50_ SARS-CoV-2 pseudovirus in 100 µL of culture medium containing 2% FBS (Gibco, Cat. No. 10099141C). Culture medium was refreshed every 12 hours. Cell culture supernatants were collected and subjected to a Gaussia luciferase assay at 48 hours post infection.

### Gaussia luciferase luminescence flash assay

To measure Gaussia luciferase expression, 20 µL of cell culture supernatant of each sample and 100 µL of assay buffer containing 4 µM coelenterazine native (Biosynth Carbosynth, Cat. No. C-7001) were added to one well of a 96-well black opaque assay plate (Corning, Cat. No. 3915), and measured with Centro LB 960 microplate luminometer (Berthold Technologies) for 0.1 second/well.

### Production and Purification of ACE2-Ig protein and SARS-CoV-2 antibodies

293F cells at the density of 6 × 10^5^ cells/mL were seeded into 100 mL SMM 293-TII serum-free medium (Sino Biological, Cat. No. M293TII) one day before transfection. Cells were then transfected with 100 µg plasmid in complex with 250 µg PEI MAX 4000 (Polysciences, Inc, Cat. No. 24765-1). Cell culture supernatants were collected at 48 to 72 hours post transfection. Human IgG1 Fc-containing proteins were purified using Protein A Sepharose CL-4B (GE Healthcare, Cat. No. 17-0780-01), eluted with 0.1 M citric acid at pH 4.5 and neutralized with 1 M Tris-HCl at pH 9.0. Buffers were then exchanged to PBS and proteins were concentrated by 30 kDa cut-off Amicon Ultra-15 Centrifugal Filter Units (Millipore, Cat. No. UFC903096).

### SARS-CoV-2 pseudovirus neutralization assay

Pseudovirus neutralization experiments were performed following our previous study^14^. In brief, SARS-CoV-2 spike variant-pseudotyped luciferase reporter viruses were pre-diluted in DMEM (2% FBS, heat-inactivated) containing titrated amounts of an ACE2-Ig construct or an anti-SARS-CoV-2 antibody. Virus-inhibitor mixtures were incubated at 37 °C for 30min, then added to HeLa-hACE2 cells in 96-well plates and incubated overnight at 37 °C. Virus-inhibitor-containing supernatant was then removed and changed with 150 µL of fresh DMEM (2% FBS) and incubated at 37 °C. Cell culture supernatants were collected for Gaussia luciferase assay at 48 h post infection.

### Biolayer interferometry (BLI) assay

The BLI assays were performed on a Fortebio Octet RED384 instrument, with the temperature and shaking speed at 30 °C and 1000 rpm respectively. ACE2-hFc constructs were diluted to 5 μg/mL in 1x assay buffer containing 150 mM NaCl, 0.1% Tween-20, 10mM HEPES and 0.1%BSA (pH 7.4), and used as ligands for the assays. RBD-mFc constructs were serially diluted to 100nM, 50nM, 25 nM, 12.5 nM, and 6.25 nM in the 1x assay buffer. Each experiment group started with a 10 min warm-up for pre-hydration of AHC biosensors, followed by cycles of baseline (60 s), loading (60 s), baseline2 (60s), association (100 s), dissociation (600 s) and regeneration plus neutralization (30 s). A 1:1 Langmuir binding model was applied for data processing. All fitted diagrams (global fit) display the entire association window and the first 200 s (or 100 s only for house mouse and Chinese rufous horseshoe bat ACE2-related assays) of dissociation phase.

### Data collection and analysis

All the experiments were independently performed for two or three times by two different people. Image Lab Software (Bio-Rad) was used to collect SDS-PAGE and Western-Blot image data. MikroWin 2000 Software (Berthold Technologies) was used to collect luciferase assay data. The BLI sensorgrams were recorded by Octet Data Acquisition 12.0 software and were fitted using Octet Data Analysis HT 12.0 software. GraphPad Prism 6.0 software was used for figure preparation and statistical analyses.

### Data availability

The study did not generate unique datasets or code. Our research resources, including methods, plasmids, and protocols, are available upon reasonable request to qualified academic investigators for noncommercial research purposes. All reagents developed in this study, such as vector plasmids, as well as detailed methods, will be made available upon written request.

## Acknowledgements

We thank the Biochemistry Core of the Shenzhen Bay Laboratory (Shenzhen, China) for providing help on performing the BLI-based binding kinetics assays and data analysis. We thank Dr. Yu J. Cao (School of Chemical Biology and Biotechnology, Peking University Shenzhen Graduate School, Shenzhen, China) for providing the 293F cells used in this study for the production of recombinant proteins. We thank Dr. Michael D. Alpert (Emmune, Inc., USA) for sharing useful comments on this manuscript.

This work was supported by Shenzhen Bay Laboratory Startup Funds (21230041, G.Z.), Major Program of Shenzhen Bay Laboratory (S201101001-2, G.Z.), Key COVID-19 Program of Shenzhen Bay Laboratory (S211410002, G.Z.), Department of Science and Technology of Guangdong province (2020B1111330001, J.Z.), and National Natural Science Foundation of China (82025001, J.Z.).

## Author contributions

G.Z. conceived and designed this study. W.Y., D.M., H.W., X.T., C.D., H.P., and Y.L. generated experimental materials. W.Y., D.M., and Y.L. performed all experiments. W.Y., D.M., Y.L., and G.Z. analyzed all data. C.L., H.L., M.F., and J.Z. contributed key resources. G.Z. and H.W. wrote the manuscript.

## References

1 Polack, F. P. et al. Safety and Efficacy of the BNT162b2 mRNA Covid-19 Vaccine. N Engl J Med 383, 2603–2615, (2020).

2 Baden, L. R. et al. Efficacy and Safety of the mRNA-1273 SARS-CoV-2 Vaccine. N Engl J Med, (2020).

3 Wang, H. et al. Development of an Inactivated Vaccine Candidate, BBIBP-CorV, with Potent Protection against SARS-CoV-2. Cell 182, 713–721 e719, (2020).

4 Xia, S. et al. Safety and immunogenicity of an inactivated SARS-CoV-2 vaccine, BBIBP-CorV: a randomised, double-blind, placebo-controlled, phase 1/2 trial. Lancet Infect Dis 21, 39–51, (2021).

5 Gao, Q. et al. Development of an inactivated vaccine candidate for SARS-CoV-2. Science 369, 77–81, (2020).

6 Logunov, D. Y. et al. Safety and immunogenicity of an rAd26 and rAd5 vector-based heterologous prime-boost COVID-19 vaccine in two formulations: two open, non-randomised phase 1/2 studies from Russia. Lancet 396, 887–897, (2020).

7 Voysey, M. et al. Safety and efficacy of the ChAdOx1 nCoV-19 vaccine (AZD1222) against SARS-CoV-2: an interim analysis of four randomised controlled trials in Brazil, South Africa, and the UK. Lancet 397, 99–111, (2021).

8 Baum, A. et al. REGN-COV2 antibodies prevent and treat SARS-CoV-2 infection in rhesus macaques and hamsters. Science 370, 1110–1115, (2020).

9 Weinreich, D. M. et al. REGN-COV2, a Neutralizing Antibody Cocktail, in Outpatients with Covid-19. N Engl J Med 384, 238–251, (2021).

10 Shi, R. et al. A human neutralizing antibody targets the receptor-binding site of SARS-CoV-2. Nature 584, 120–124, (2020).

11 Jones, B. E. et al. The neutralizing antibody, LY-CoV555, protects against SARS-CoV-2 infection in non-human primates. Sci Transl Med, (2021).

12 Gottlieb, R. L. et al. Effect of Bamlanivimab as Monotherapy or in Combination With Etesevimab on Viral Load in Patients With Mild to Moderate COVID-19: A Randomized Clinical Trial. JAMA 325, 632–644, (2021).

13 Zhou, P. et al. A pneumonia outbreak associated with a new coronavirus of probable bat origin. Nature 579, 270–273, (2020).

14 Li, Y. et al. SARS-CoV-2 and Three Related Coronaviruses Utilize Multiple ACE2 Orthologs and Are Potently Blocked by an Improved ACE2-Ig. J Virol 94, (2020).

15 Shi, J. et al. Susceptibility of ferrets, cats, dogs, and other domesticated animals to SARS-coronavirus 2. Science 368, 1016–1020, (2020).

16 Cui, J., Li, F. & Shi, Z. L. Origin and evolution of pathogenic coronaviruses. Nat Rev Microbiol 17, 181–192, (2019).

17 Li, F. Receptor recognition and cross-species infections of SARS coronavirus. Antiviral Res 100, 246–254, (2013).

18 Letko, M., Marzi, A. & Munster, V. Functional assessment of cell entry and receptor usage for SARS-CoV-2 and other lineage B betacoronaviruses. Nat Microbiol 5, 562–569, (2020).

19 Hoffmann, M. et al. SARS-CoV-2 Cell Entry Depends on ACE2 and TMPRSS2 and Is Blocked by a Clinically Proven Protease Inhibitor. Cell, (2020).

20 Wang, J. et al. Mouse-adapted SARS-CoV-2 replicates efficiently in the upper and lower respiratory tract of BALB/c and C57BL/6J mice. Protein Cell 11, 776–782, (2020).

21 Dinnon, K. H., 3rd et al. A mouse-adapted model of SARS-CoV-2 to test COVID-19 countermeasures. Nature 586, 560–566, (2020).

22 Gu, H. et al. Adaptation of SARS-CoV-2 in BALB/c mice for testing vaccine efficacy. Science 369, 1603–1607, (2020).

23 Li, Y. et al. Single amino-acid change in mouse Ace2 or viral spike enables SARS-CoV-2 to utilize mouse Ace2. (2021. manuscript submitted).

24 Kaushal, N. et al. Mutational Frequencies of SARS-CoV-2 Genome during the Beginning Months of the Outbreak in USA. Pathogens 9, (2020).

25 Gribble, J. et al. The coronavirus proofreading exoribonuclease mediates extensive viral recombination. PLoS Pathog 17, e1009226, (2021).

26 Thomson, E. C. et al. The circulating SARS-CoV-2 spike variant N439K maintains fitness while evading antibody-mediated immunity. bioRxiv (2020). doi: https://doi.org/10.1101/2020.11.04.355842

27 Davies, N. G. et al. Estimated transmissibility and severity of novel SARS-CoV-2 Variant of Concern 202012/01 in England. medRxiv (2020). doi:10.1101/2020.12.24.20248822

28 Laydon, D. J. et al. Transmission of SARS-CoV-2 Lineage B.1.1.7 in England: Insights from linking epidemiological and genetic data. MedRxiv (2020). doi: https://doi.org/10.1101/2020.12.30.20249034

29 Tegally, H. et al. Emergence and rapid spread of a new severe acute respiratory syndrome-related coronavirus 2 (SARS-CoV-2) lineage with multiple spike mutations in South Africa. medRxiv, (2020).

30 Diseases, J. N. I. o. I. Brief report: New Variant Strain of SARS-CoV-2 Identified in Travelers from Brazil. (2021). doi:https://www.niid.go.jp/niid/images/epi/corona/covid19-33-en-210112

31 Elaswad, A., Fawzy, M., Basiouni, S. & Shehata, A. A. Mutational spectra of SARS-CoV-2 isolated from animals. PeerJ 8, e10609, (2020).

32 Oude Munnink, B. B. et al. Transmission of SARS-CoV-2 on mink farms between humans and mink and back to humans. Science 371, 172–177, (2021).

33 Kemp, S. A. et al. Neutralising antibodies drive Spike mediated SARS-CoV-2 evasion. MedRxiv, (2020).

34 Li, Q. et al. The Impact of Mutations in SARS-CoV-2 Spike on Viral Infectivity and Antigenicity. Cell 182, 1284–1294 e1289, (2020).

35 Peacock, T. P., Penrice-Randal, R., Hiscox, J. A. & Barclay, W. S. SARS-CoV-2 one year on: evidence for ongoing viral adaptation. J Gen Virol 102, (2021).

36 Chen, L. et al. RNA based mNGS approach identifies a novel human coronavirus from two individual pneumonia cases in 2019 Wuhan outbreak. Emerg Microbes Infect 9, 313–319, (2020).

37 Thomson, E. C. et al. Circulating SARS-CoV-2 spike N439K variants maintain fitness while evading antibody-mediated immunity. Cell 184, 1171–1187 e1120, (2021).

38 Welkers, M. R. A., Han, A. X., Reusken, C.B.E.M. & Eggink, D. Possible host-adaptation of SARS-CoV-2 due to improved ACE2 receptor binding in mink. Virus evolution doi:10.1093/ve/veaa094, (2020).

39 Volz, E. et al. Assessing transmissibility of SARS-CoV-2 lineage B.1.1.7 in England. Nature 593, 266–269, (2021).

40 Davies, N. G. et al. Estimated transmissibility and impact of SARS-CoV-2 lineage B.1.1.7 in England. Science 372, (2021).

41 Washington, N. L. et al. Emergence and rapid transmission of SARS-CoV-2 B.1.1.7 in the United States. Cell 184, 2587–2594 e2587, (2021).

42 Motozono, C. et al. An emerging SARS-CoV-2 mutant evading cellular immunity and increasing viral infectivity. bioRxiv, (2021).

43 Cai, Y. et al. Structural basis for enhanced infectivity and immune evasion of SARS-CoV-2 variants. bioRxiv, (2021).

44 Gobeil, S. M. et al. Effect of natural mutations of SARS-CoV-2 on spike structure, conformation and antigenicity. bioRxiv, (2021).

45 Zhu, X. et al. Cryo-electron microscopy structures of the N501Y SARS-CoV-2 spike protein in complex with ACE2 and 2 potent neutralizing antibodies. PLoS Biol 19, e3001237, (2021).

46 Chan, K. K. et al. Engineering human ACE2 to optimize binding to the spike protein of SARS coronavirus 2. Science 369, 1261–1265, (2020).

47 Glasgow, A. et al. Engineered ACE2 receptor traps potently neutralize SARS-CoV-2. Proc Natl Acad Sci U S A 117, 28046–28055, (2020).

48 Montagutelli, X. et al. The B1.351 and P.1 variants extend SARS-CoV-2 host range to mice. BioRxiv, (2021).

49 Ulrich, L., Wernike, K., Hoffmann, D., Mettenleiter, T. C. & Beer, M. Experimental Infection of Cattle with SARS-CoV-2. Emerg Infect Dis 26, 2979–2981, (2020).

50 Falkenberg, S. et al. Experimental Inoculation of Young Calves with SARS-CoV-2. Viruses 13, (2021).

51 Pickering, B. S. et al. Susceptibility of Domestic Swine to Experimental Infection with Severe Acute Respiratory Syndrome Coronavirus 2. Emerg Infect Dis 27, 104–112, (2021).

52 Meekins, D. A. et al. Susceptibility of swine cells and domestic pigs to SARS-CoV-2. Emerg Microbes Infect 9, 2278–2288, (2020).

53 Wu, K. et al. mRNA-1273 vaccine induces neutralizing antibodies against spike mutants from global SARS-CoV-2 variants. bioRxiv, (2021).

54 Wang, Z. et al. mRNA vaccine-elicited antibodies to SARS-CoV-2 and circulating variants. bioRxiv, (2021).

55 Rees-Spear, C. et al. The impact of Spike mutations on SARS-1 CoV-2 neutralization. bioRxiv, (2021).

56 Zhou, D. et al. Evidence of escape of SARS-CoV-2 variant B.1.351 from natural and vaccine-induced sera. Cell 184, 2348–2361 e2346, (2021).

57 Wang, P. et al. Antibody resistance of SARS-CoV-2 variants B.1.351 and B.1.1.7. Nature 593, 130–135, (2021).

58 Shinde, V. et al. Efficacy of NVX-CoV2373 Covid-19 Vaccine against the B.1.351 Variant. N Engl J Med, (2021).

59 Li, W. et al. Angiotensin-converting enzyme 2 is a functional receptor for the SARS coronavirus. Nature 426, 450–454, (2003).

60 Gardner, M. R. et al. AAV-expressed eCD4-Ig provides durable protection from multiple SHIV challenges. Nature 519, 87–91, (2015).

61 Wu, X. et al. Rational design of envelope identifies broadly neutralizing human monoclonal antibodies to HIV-1. Science 329, 856–861, (2010).

62 Brass, A. L. et al. The IFITM proteins mediate cellular resistance to influenza A H1N1 virus, West Nile virus, and dengue virus. Cell 139, 1243–1254, (2009).

63 Bailey, C. C., Huang, I. C., Kam, C. & Farzan, M. Ifitm3 limits the severity of acute influenza in mice. PLoS Pathog 8, e1002909, (2012).

64 Huang, I. C. et al. Distinct patterns of IFITM-mediated restriction of filoviruses, SARS coronavirus, and influenza A virus. PLoS Pathog 7, e1001258, (2011).

65 Moore, M. J. et al. Retroviruses pseudotyped with the severe acute respiratory syndrome coronavirus spike protein efficiently infect cells expressing angiotensin-converting enzyme 2. J Virol 78, 10628–10635, (2004).

66 Ren, W. et al. Difference in receptor usage between severe acute respiratory syndrome (SARS) coronavirus and SARS-like coronavirus of bat origin. J Virol 82, 1899–1907, (2008).

67 Lei, C. et al. Neutralization of SARS-CoV-2 spike pseudotyped virus by recombinant ACE2-Ig. Nat Commun 11, 2070, (2020).

68 Li, Q. et al. SARS-CoV-2 501Y.V2 variants lack higher infectivity but do have immune escape. Cell 184, 2362–2371 e2369, (2021).

69 World Health Organization. Laboratory procedures: serological detection of avian influenza A(H7N9) infections by microneutralization assay. May 2013. https://www.who.int/influenza/gisrs_laboratory/cnic_serological_diagnosis_microneutralization_a_h7n9.pdf

